# Adaptive Safety Coding in the Prefrontal Cortex

**DOI:** 10.1101/2024.07.19.604228

**Authors:** Sarah M. Tashjian, Joseph Cussen, Wenning Deng, Bo Zhang, Dean Mobbs

## Abstract

Pivotal to self-preservation is the ability to identify when we are safe and when we are in danger. Previous studies have focused on safety estimations based on the features of external threats and do not consider how the brain integrates other key factors, including estimates about our ability to protect ourselves. Here we examine the neural systems underlying the online dynamic encoding of safety. The current preregistered study used two novel tasks to test four facets of safety estimation: *Safety Prediction*, *Meta-representation*, *Recognition*, and *Value Updating*. We experimentally manipulated safety estimation changing both levels of external threats and self-protection. Data were collected in two independent samples (behavioral *N*=100; fMRI *N*=30). We found consistent evidence of subjective changes in the sensitivity to safety conferred through protection. Neural responses in the ventromedial prefrontal cortex (vmPFC) tracked increases in safety during all safety estimation facets, with specific tuning to protection. Further, informational connectivity analyses revealed distinct hubs of safety coding in the posterior and anterior vmPFC for external threats and protection, respectively. These findings reveal a central role of the vmPFC for coding safety.

---

In their natural habitat, the Sundarbans Tiger, which is a formidable threat to humans, is justifiably feared. However, with a gun in hand, we fear the Tiger less. At the zoo, behind the protection of laminated glass, fear is replaced with enthused curiosity. In all of these scenarios, the sensory features of the Tiger remain stable, yet the perception of safety fluctuates. These fluctuations in safety occur as a function of information unrelated to the Tiger’s features, but changes in the perception of safety. Despite this knowledge, the contributions of safety estimation beyond external threats are largely ignored in the existing literature. To accurately estimate safety, we need to integrate threat-related information with information about our ability to protect ourselves.^1,2^ From an evolutionary perspective, these factors determine how likely we are to succeed in surviving encounters with natural dangers.

The ability to recognize safety is critical to adjusting adaptive defensive responses, reducing stress, and initiating in other survival behaviors, including foraging and mating.^3,4^ How the brain integrates multiple sources of dynamic information to compute safety estimates remains to be identified. One theory is that the coding of safety is constructed based on representations that reflect the learned and phenotypic features of external threats.^5–7^ The brain creates an internal model that integrates incoming sensory information about a stimulus with other relevant information such as context (i.e., Am I at the zoo or alone in the forest?). This ‘other’ information does not pertain to the threat itself but is important for accurately estimating safety. As relevant information changes so can the safety estimate even if the threat itself remains unchanged. Similarly, changing threat features (i.e., Is the tiger awake or asleep?) should trigger updated safety estimates through integration in safety circuits with consequences for survival behavior. Such *meta-representations* are likely to involve neuronal ensembles that integrate the valuation of threats with information about our ability to protect against them.^8,9^ Depending on the resulting calculation, defensive excitation or inhibition occurs.^10–13^

We test two primary hypotheses regarding the functioning of safety neural circuitry. First, we test the hypothesis that the neural systems involved in representing threat and protection are dissociable during *Safety Prediction*.^2^ We extend beyond models focused on the external environment to examine fluctuations in safety as they relate to self-relevant states (e.g., the value of protection). We argue that self-relevant states – the extent to which one can successfully protect oneself – are a crucial, but overlooked, factor in estimating safety. Going face-to-face with a tiger with no weapon in hand will likely result in death. Yet, with a knife, we have an increased chance of deterring the predator and surviving. A gun further increases survival likelihood and turns the tables on the tiger as the likely casualty. Second, we hypothesize that the brain integrates threat and protective information to confer a safety ‘*meta-representation’*.^14–16^ To test this, we manipulated the safety value of threat and protective stimuli as a function of their pairing without altering the perceptual features of either stimulus. For example, a tiger becomes less dangerous when faced with a gun as opposed to a knife, but the tiger itself does not change. We hypothesize that the safety modulator (e.g., the gun or knife) will be *meta-represented* during the evaluation of the modulated stimulus (e.g., the tiger), even though the modulator is not being displayed and therefore not visually perceived.

We hypothesize a candidate region for human safety coding is the ventromedial prefrontal cortex (vmPFC). Recent theoretical developments suggest representations of threat and protective information are encoded in canonical defensive and cognitive neural circuits, the latter including the vmPFC.^2,17–22^ We propose that threats are computed in a bottom-up fashion, primarily driven by sensory processing regions of the brain, whereas protection is integrated with threat information through top-down metacognitive circuitry related to self-evaluation.^23,24^ Research on threat controllability provides support for this supposition - controllability improves fear extinction in humans implying a role of the self in safety estimation.^25^ Work outside of the threat context points to the vmPFC as pivotal in supporting decisions concerning the self.^16^ The vmPFC aids the affective processing of safety signals as well as the acquisition of new threat associations and threat extinction during Pavlovian threat conditioning.^2,17–22,26–31^ Thus, the vmPFC is an important region of interest for testing how the brain integrates information to formulate safety estimations.

Although we propose that successful safety estimation in humans relies on integrating multiple distinct components, how the brain interprets and weighs these factors remains unknown. No prior study has systematically manipulated both threat and protection concerning safety judgments. In two preregistered samples using a novel Safety Estimation Task (Fig. 1a) (*N*=100 behavioral; *N*=30 functional magnetic resonance imaging, fMRI; https://osf.io/hw3r9), we examined how subjects evaluated different types of safety information (*Safety Prediction*), as well as how they integrated information to estimate safety when perceptually identical stimuli changed in value (*Safety Meta-representation*). As a comparison, we examined neural activation when safety was certain during the outcome phase (*Safety Recognition*). We also tested how safety estimation changed as a function of experience (*Safety Value Updating*) in a separate task administered before and after the Safety Estimation Task. We examined safety coding at the whole-brain level but focused on the contributions of the vmPFC as a hypothesized safety coding hub.

**Figure 1.**
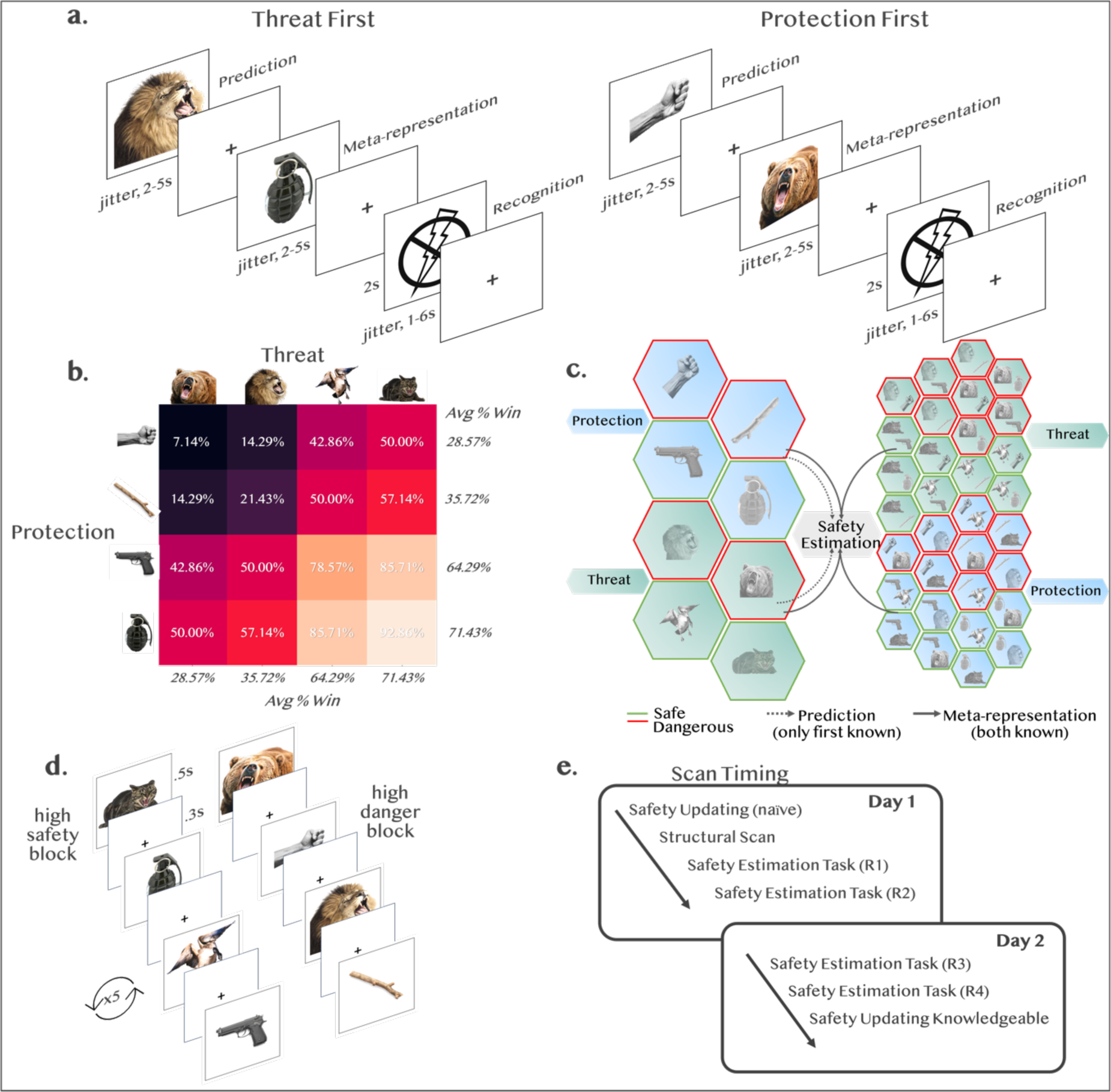
(a) Safety Estimation Task. Subjects saw stimuli pairs comprised of a threat (animal) and protection (weapon) with presentation of threat/protection counterbalanced. First a weapon or animal was presented (*Safety Prediction*) and subjects made an initial estimation of whether they would win or lose the battle. Then the paired stimulus was presented (*Safety Meta-representation*) and subjects made an updated judgment as to whether they would win or lose. After both stimuli were presented, subjects saw the outcome of the battle depicted as either a shock (loss) or no shock (win) (*Safety Recognition*). **(b) Safety probabilities.** Stimuli along each of the threat and protection continuum had equivalent shock probabilities. Italics depict average shock value for each stimulus irrespective of pairing. Paired probabilities are depicted in the heatmap. **(c) Model of safety estimation.** When only the first stimulus is known, *Prediction* (left) is biased and based only on the potency and protective value of the first stimulus presented, with variability depending on relevance (i.e., whether it is an external threat or self-relevant protection). When the second stimulus is presented, *Meta-representations* (right) integrate information about the first and second stimulus to update safety estimates accounting for the contribution of each stimulus. Both phases converge on a final estimate of safety. **(d) Safety Value Updating Task.** Blocks of stimuli were presented according to safety value with each stimulus presented 5x in a block. Subjects were told to look at the stimuli carefully and that no judgments were to be made and no shocks would be delivered. **(e) Scan session timing.** On the first day subjects completed the Naive Safety Value Updating Task, a structural scan, and the first and second runs of the Safety Estimation Task. On the second day, subjects completed the third and fourth runs of the Safety Estimation Task followed by the Knowledgeable Safety Value Updating Task.

During the Safety Estimation Task, subjects were shown stimuli pairs comprised of an external threat (dangerous animal) and a self-relevant protection (powerful weapon). Four stimuli for each threat and protection were used (Fig. 1b). Presentation of stimuli was counterbalanced such that in some trials the protection was shown first and in other trials, the protection was shown second. Subjects made binary forced-choice judgments about whether they thought they would win or lose the battle against the dangerous animal using the weapon they were provided. All trials included a threat and protection, separately presented to allow for analyzing subject response when shown the first stimulus (*Safety Prediction*) separate from the second stimulus (*Safety Meta-representation*). Safety probabilities for threat and protection stimuli were matched such that high safe protection and high safe threat were equally likely to result in a win. Combinations of threat and protection stimuli were also matched with safety outcomes varying as a function of the average safety value of each stimulus in the pair. After both stimuli were presented, subjects saw the outcome of the battle (*Safety Recognition*). Lost battles risked delivery of an electric shock to the subject’s wrist (randomly delivered on 20% of lost trials). Successful battles resulted in 100% safety with no electric shock.

*Safety Value Updating* was tested in a separate, passive viewing task administered before and after the Safety Estimation Task (Fig. 1d-e). All stimuli from the Safety Estimation Task were presented in blocks of stimuli subsets depending on the safety value of each stimulus (e.g., the high safety block consisted of the weapons and animals with the highest safety probability). The first “naive” viewing was performed while subjects had no information about the Safety Estimation Task. The second “knowledgeable” viewing was performed after all runs of the Safety Estimation Task were completed and subjects had experienced the stimuli as relevant to their safety. Neural regions activated at the second viewing, but not the first viewing, were interpreted as tracking updated safety values. Dangerous weapons were used as protection during the Safety Estimation Task, meaning at the naive viewing all stimuli appeared dangerous, but, at the knowledgeable viewing, weapons with high safety value should be updated to indicate protection.

## Results

Results are reported as facets of safety estimation, (1) *Safety Prediction* in response to the first stimulus presentation during the Safety Estimation Task when subjects could estimate safety based only on partial information, (2) *Safety Meta-representation* at the second stimulus presentation during the Safety Estimation Task when subjects could estimate safety on full information (Fig. 1c), (3) *Safety Recognition* at the outcome during the Safety Estimation Task when subjects knew whether they were at risk of electric shock, and (4) *Safety Value Updating* comparing the “knowledgeable” and “naive” rounds of the passive viewing task. For each section, behavioral models are reported as well as univariate and multivariate fMRI results. Safety value (high versus low) and safety relevance (self versus external) are both considered.

### Safety Prediction

Differences in *Safety Prediction* represented a bias toward initial safety information as a function of relevance (external versus self), given that safety probabilities for threat and protection were identical.

Behaviorally, subjects in both the behavioral and MRI samples tracked probabilities of winning and losing across the safety continuum such that subjects estimated a higher probability of winning for stimuli with higher safety probabilities (Fig. 2a; Table S1). Safety relevance (external versus self) affected initial safety bias such that subjects estimated greater safety when the first stimulus presented was protection, behavioral sample difference between protection and threat *α*=.23, 95%CI[.21, .25]; MRI sample *α*=.28, 95%CI[.25, .31] (Fig. 2b).

**Figure 2.**
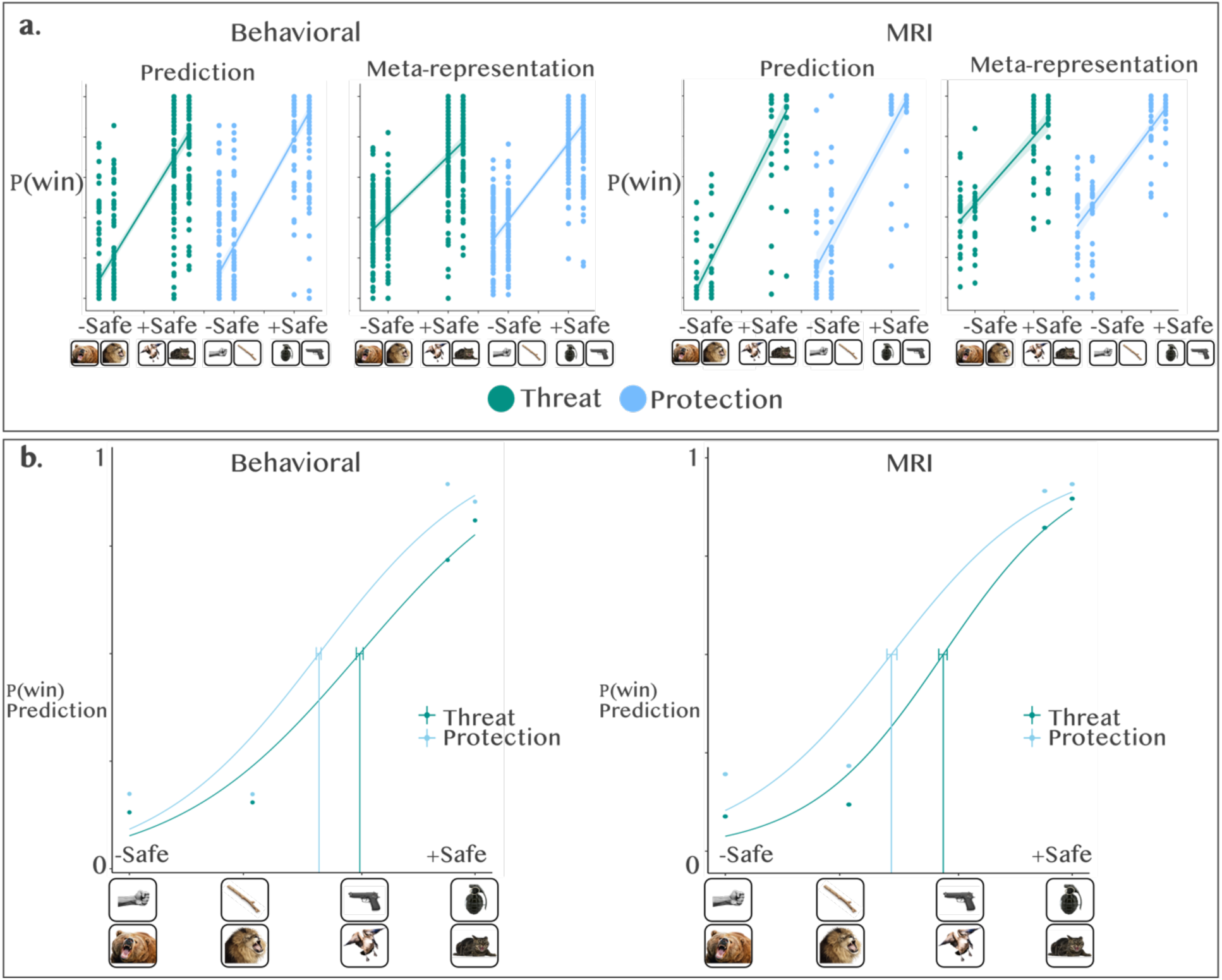
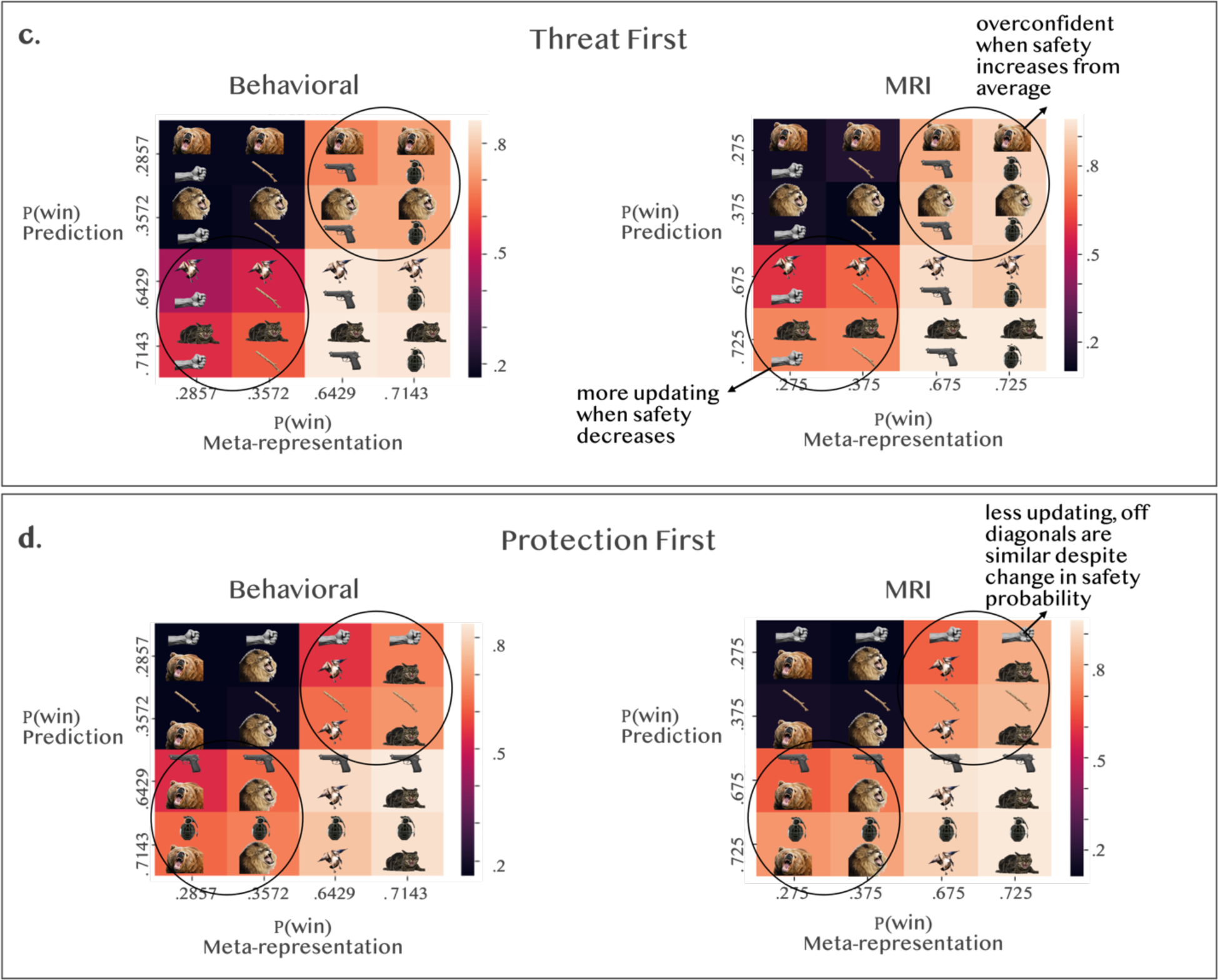
Behavioral results. Subjects tracked safety probabilities across the safety continuum for both stimulus types. **(a)** Linear regression depicting the association between objective safety and subjective safety estimate for threat and protection stimuli. Mixed effects model results indicated subjects differentiated safety in accordance with the objective safety continuum. See Supplemental Table S1. x-axis=safety continuum, y-axis= subjective probability of winning (0=lose, 1=win). **(b)** Psychometric curves for *Safety Prediction* in both samples. Subjects reached the safety detection threshold faster when protection was presented as the first stimulus in the battle pair. behavioral *N=100* (left), MRI *N*=30 (right). **(c)** When threatening animals were the first stimulus, subjects were more likely to update *Safety Meta-representations* in response to protection, particularly when safety decreased from the average. **(d)** When protection was the first stimulus, subjects were more likely to meta-represent the first stimulus and not update the safety estimates as indicated by similar off diagonals despite safety probabilities shifting from average in opposite directions. Panels c and d indicate subjects updated *Safety Meta-representations* to a greater extent when protection information conflicted with initial threat information but not when threat information conflicted with initial protection information. x-axis values: safety change 1 = high safety probability followed by high danger probability (e.g., gun followed by a grizzly; cat followed by stick); safety change 2 = high danger probability followed by a high safety probability (e.g., grizzly followed by grenade; stick followed by a cat).

Univariate fMRI analyses showed that the brain tracked safety value in response to the first stimulus presented, with dissociable circuits activated depending on safety value. As the stimuli increased in safety value, so did activation in the vmPFC, as hypothesized (Fig. 3a, Table S2). Considering each stimulus type separately, increasing safety for threatening animals activated the lateral occipital cortex (Fig. 3b), and increasing safety for protection activated the vmPFC, amygdala, temporal pole, and anterior cingulate cortex (ACC) (Fig. 3c).

**Figure 3.**
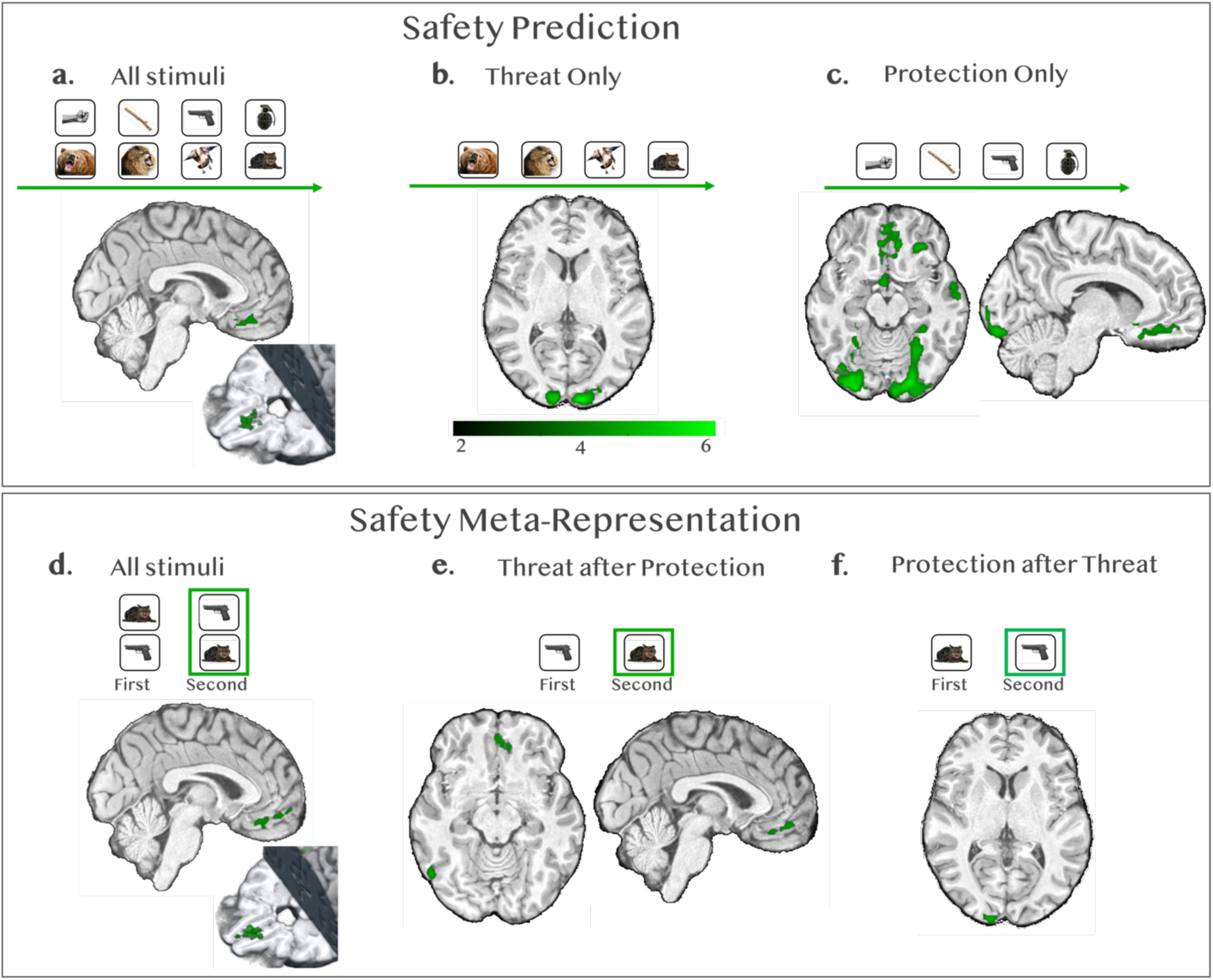
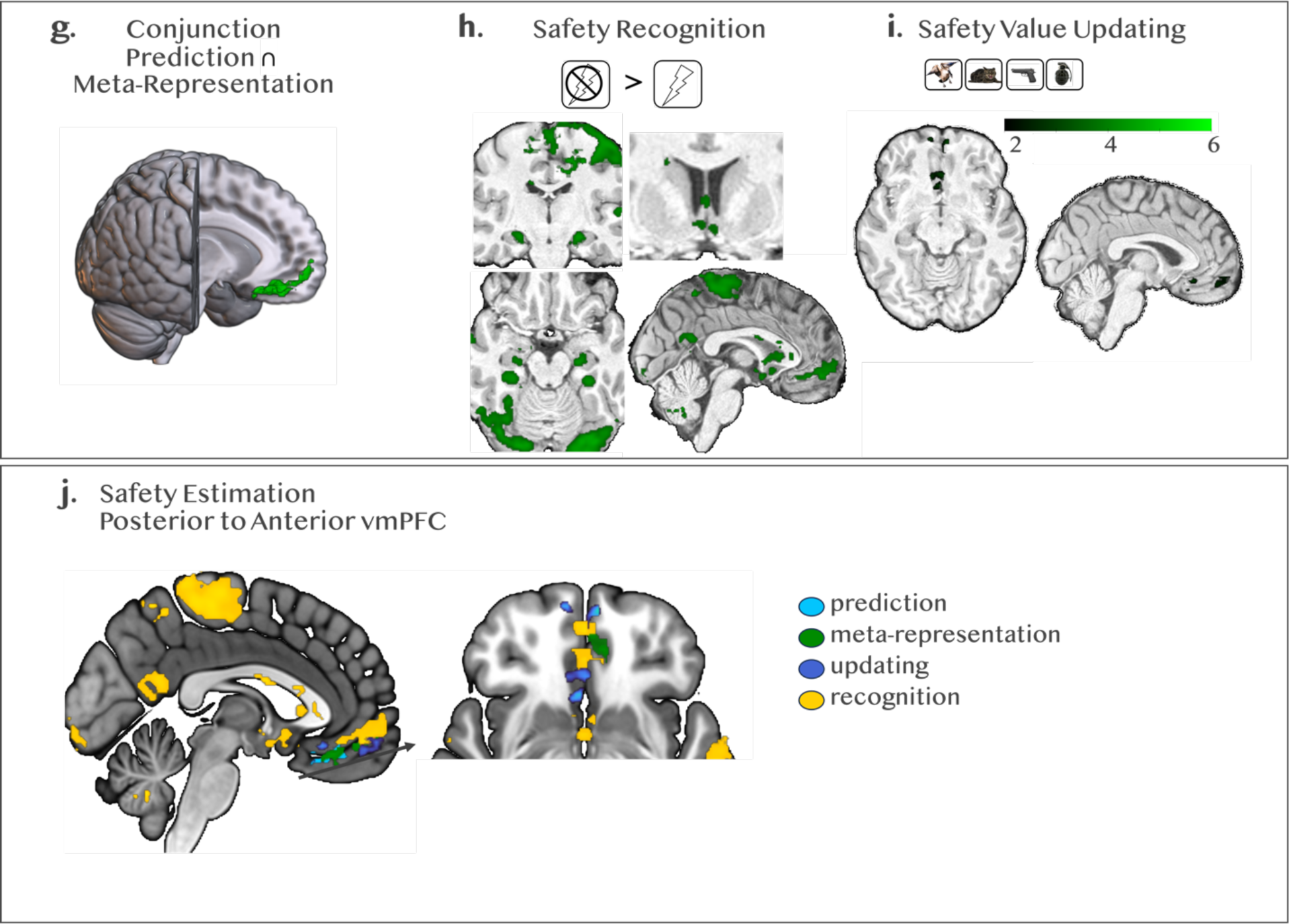
Neural Activation. Parametric increases in whole-brain neural activity that track the increased objective safety value of stimuli. Analyses in (3a-e, g) were conducted using FSL Randomise, TFCE, FWE-corrected *p*<.05. Color bar indicates t-intensity values. **(a-c) Safety Prediction.** During the Safety Estimation Task, the first stimulus presented represented a bias to partial information. Activation increased as safety increased based on the average safety probability of each stimulus. Significant clusters indicate activation increased in those regions in response to stimuli as they increased in safety on the shock probability continuum (e.g., greater response to a grenade than a fist). (a) Threat and Protection collapsed, (b) Threat only, (c) Protection only. **(d-f) Safety Meta-representation.** During the second stimulus presentation, safety value was determined by integrating information about the first stimulus with the second stimulus. Safety was based on comparison with the average safety value of the stimulus. For example, if a stick was shown as the second stimulus and was paired with a lion, the probability of shock would increase from 64.28% on average to 78.57%, but when paired with the cat the stick probability of shock would decrease to 42.86. The stick after the cat, but not the lion would be considered an ‘increase’ in safety. (d) Threat and Protection collapsed, (e) Threat only, (f) Protection only. **(g) Conjunction Prediction** ∩ **Meta-representation.** Conjunction analyses for the safety prediction increasing safety > increasing danger ∩ safety meta-representation increasing safety > increasing danger for all stimuli, with overlapping activation in the vmPFC; *Z=2.3, p<.05*. **(h) Safety Recognition.** Neural activation in response to successful battles during the outcome screen indicating 100% certainty of safety compared with unsuccessful battles during the outcome screen that indicated potential for electric shock. **(i) Safety Value Updating.** Multivariate searchlight revealed neural activation change in the vmPFC when subjects viewed stimuli with high safety value, contrast of Knowledge > Naive. Searchlight was *a priori* restricted to the vmPFC using an ROI defined via neurovault. **(j) Safety Estimation.** vmPFC overlap for all stages of safety estimation demonstrate posterior to anterior shift as safety becomes more certain.

Activation corresponding to decreases in safety was observed in the occipital pole and postcentral gyrus (all stimuli and threatening animals, Supplemental Fig. S1a-b). Decreasing the safety value of protection did not show significant differences in parametric activation (Supplemental Fig. S1c).

Informational Connectivity (IC) testing multi-voxel pattern synchronization between regions of interest (ROIs, Fig. 4a) revealed a dense network involved in coding *Safety Prediction* (Fig. 4b). The anterior cingulate cortex (ACC, caudodorsal), insula (anterior), striatum (dorsal and ventral), vmPFC (anterior and posterior), thalamus, and hippocampus were connected during safety decoding. Betweenness Centrality (BC) revealed the ACC as the primary hub of the network, with the insula, dorsal striatum, and posterior vmPFC also showing centrality contributions. While encoding the safety value of protective weapons, the anterior vmPFC emerged as the central hub, and the thalamus was eliminated as part of the network. While encoding safety in response to threatening animals, the posterior vmPFC emerged as the network hub and connections with the hippocampus, thalamus, and ventral striatum were no longer significant.

**Figure 4.**
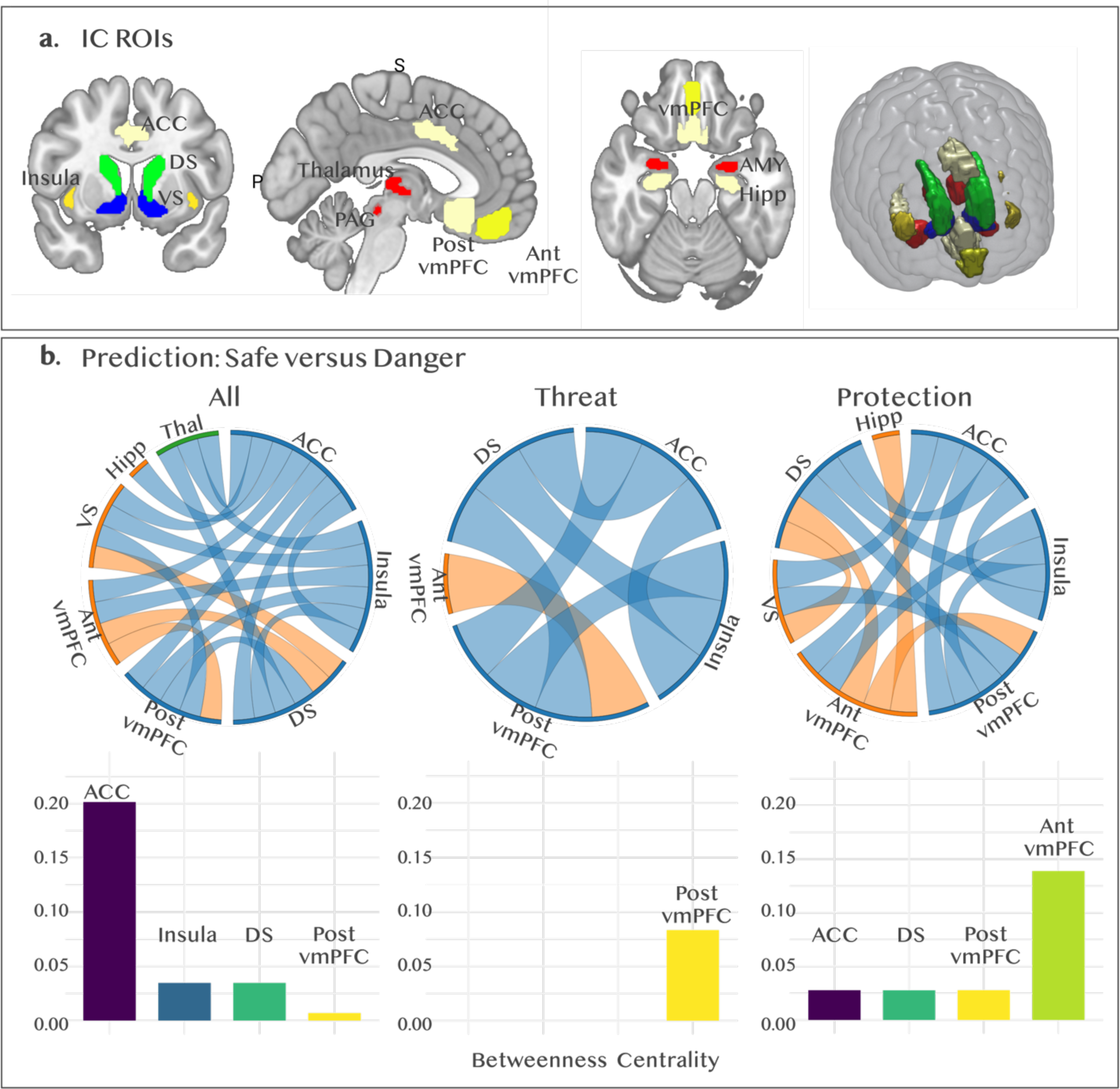
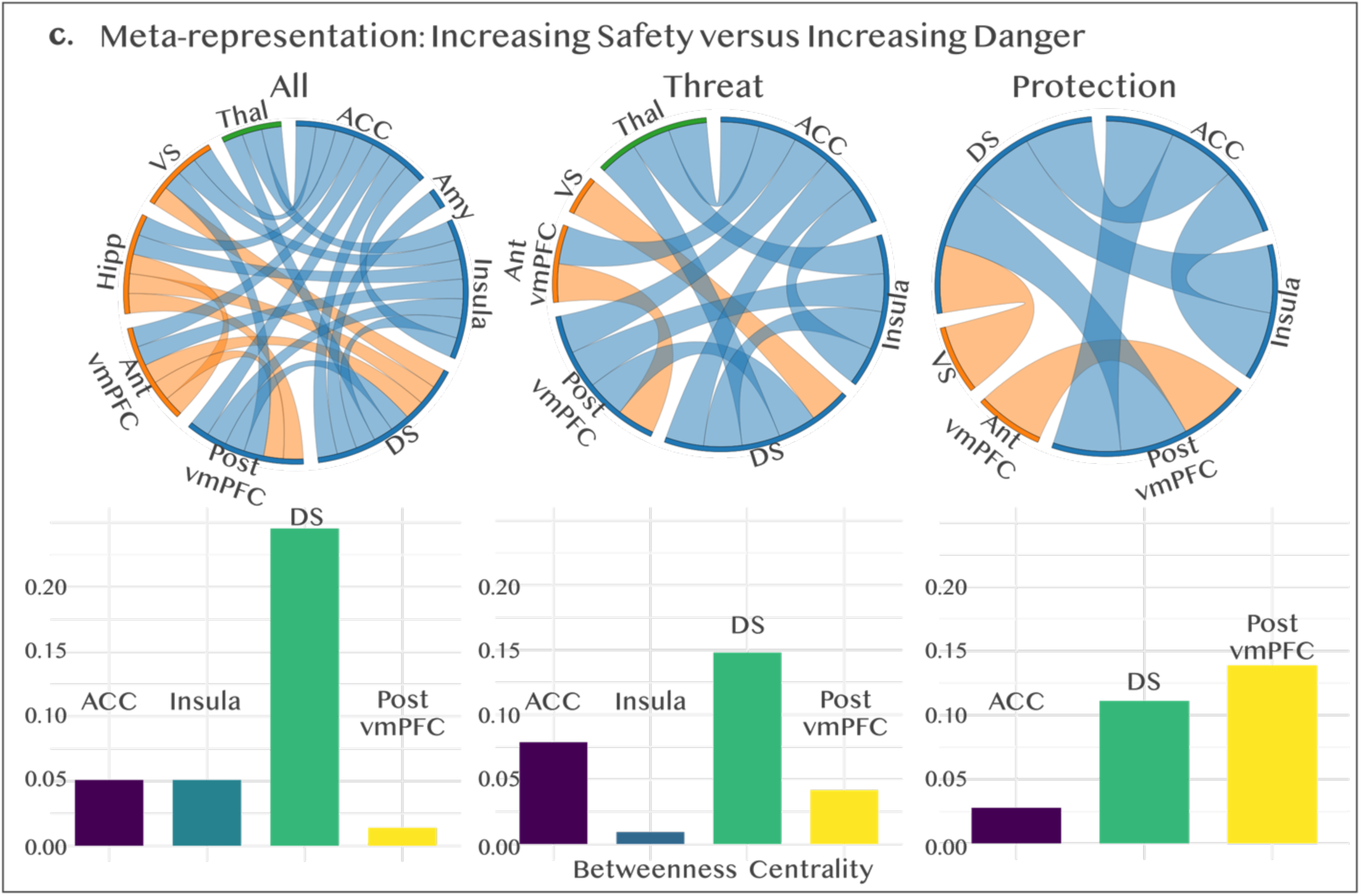
Safety Network Connectivity. Multivariate functional connectivity (Informational Connectivity) between **(a)** ten *a priori* selected ROIs. Informational connectivity was computed by using covariation trial-by-trial decoding accuracy between a pair of regions. **(b)** This resulted in a connectivity matrix between ROIs for the first stimulus presentation indicating regions that communicated while decoding states of safety for all stimuli collapsed, threat stimuli, and protection stimuli (left to right), all connections *p*<0.05. We computed betweenness centrality (BC) within this network to find hubs connecting regions during decoding. **(c)** For second stimulus presentation, regions of connectivity were decoded based on whether stimuli increased in safety or increased in danger as a function of the stimulus pairing), all connections *p*<0.05. For example, a lion paired with a fist would increase in danger whereas a lion paired with a grenade would decrease in danger.

### Safety Meta-representation

During *Safety Meta-representation*, stimuli were examined in terms of safety fluctuation: when the same stimulus had a higher-than-average safety value (e.g., a fist paired with a cat) compared with a lower-than-average safety value (e.g., a fist paired with a grizzly). Fluctuation in neural response reflects the integration of information about the initial stimulus encountered (e.g., cat, grizzly) with the second stimulus presented (e.g., fist) while holding the perceptual experience of the second stimulus constant.

Subjects’ safety meta-representation behavior was consistent with their predictive behavior such that subjects in both samples estimated a higher probability of winning for stimuli with higher safety probabilities (Fig. 2a; Table S1). There was no significant difference in the threshold of safety detection as a function of the second stimulus type, behavioral sample difference between protection and threat *α*=.01, 95%CI[-.03, .04]; MRI sample *α*=-.02, 95%CI[-.09, .04].

During *Safety Meta-representation*, subjects fixated more on the initial safety value if protection stimuli were presented first, behavioral *t*(99)=2.93, *p*=.004, Cohen’s *d*=.29, *α*_protection_=.50, *SD*_protection_=.22, *α*_threat_=.39, *SD*_threat_=.20; MRI *t*(29)=.33, *p*=.75, Cohen’s *d*=.06, *α*_protection_=.47, *SD*_protection_=.21, *α*_threat_=.44, *SD*_threat_=.18. In other words, when threats were presented first, subjects updated their safety estimations to a greater extent in response to protection information if that information conflicted with the initial average estimate conferred by the threat stimulus (Fig. 2c). Subjects rated winning probabilities as higher if the second stimulus was a powerful weapon following a dangerous animal, compared to a weak weapon following a safe animal (75% versus 53%), despite equivalent objective probabilities (Fig. 2c). However, subjects rated safety probabilities as equivalent when a dangerous animal followed a weak weapon or a safe animal followed a powerful weapon (63% for both) (Fig. 2d).

When stimuli increased in safety value compared to its average, vmPFC activation increased parametrically (Fig. 3d, Table S2), evincing a more distributed pattern of activation across the vmPFC compared with *Safety Prediction*. Considering each stimulus type separately, neural response to external threats increased in safety (as a function of being paired with more powerful weapons) activation increased in the vmPFC and lateral occipital cortex (Fig. 3e). As self-relevant protective stimuli increased safety, neural response increased in the occipital cortex (Fig. 3f). Conjunction analyses identified the vmPFC as a common neural substrate of *Safety Prediction* and *Meta-representation* (Fig. 3g).

In response to increasing danger, activation increased in the insula, thalamus, anterior cingulate cortex, and PAG (Supplemental Fig. S1d), showing a more typical pattern of defensive circuitry activation than that observed in response to dangerous stimuli at first presentation. As threatening animals become less safe, the bilateral thalamus, right insula, pre-supplementary motor cortex, and medial occipital cortex parametrically responded with increased activation (Supplemental Fig. S1e). When self-protective stimuli were rendered less safe than average because of the initial danger value of the threat, neural response increased in the insula, thalamus, and ACC (Supplemental Fig. S1f).

Testing multi-voxel pattern synchronization using IC, the same safety network was decoded with the addition of the amygdala in response to *Safety Meta-representation* (Fig. 4c). IC during *Safety Meta-representation* tracked changes in safety from overall average values, in line with univariate results. BC revealed a switch from the ACC as the top *Safety Prediction* hub to the dorsal striatum as the top *Safety Meta-representation* hub when examining all stimuli and in response to threatening animals, with the ACC, insula, and posterior vmPFC playing centrality roles. The posterior vmPFC was the central hub for the *meta-representation* of protective weapons, followed by the dorsal striatum and ACC and the insula no longer showing as a central hub.

### Safety Recognition

*Safety Recognition* was tested in response to the trial outcome when subjects won the battle and were safe from electric shock. Activation in the vmPFC as well as in the striatum and bilateral hippocampus increased in response to safe outcomes during the task (Fig. 3h).

In response to negative outcomes (lost battles conferring risk of shock), the bilateral insula and PAG were activated (Supplemental Fig. S1g).

### Safety Value Updating

*Safety Value Updating* was examined during a separate fMRI task. Subjects viewed all threat and protection stimuli in a rapid block design before the Safety Estimation Task (naive first viewing) and then again after performing the full Safety Estimation Task (knowledgeable second viewing). This allowed us to test whether subjects updated their response to stimuli with high safety values after learning about safety probabilities during the Safety Estimation Task. After *Safety Value Updating*, targeted multivariate searchlight revealed significant changes in vmPFC representation of stimuli with high safety probabilities (cat, goose, gun grenade) (Fig. 3i). No significant changes from pre- to post-task emerged for stimuli with high danger probabilities (lion, grizzly, fist, stick).

### vmPFC

Results show vmPFC involvement at all stages of safety estimation (Fig. 3j) with posterior anterior gradients as safety increases in certainty from *Prediction* (partial information) to *Meta-representation* (full information, outcome unknown) to *Value Updating* (no outcome relevance, stimulus value learned) to *Recognition* (full certainty of safety).

## Discussion

This study identifies neural systems involved in safety coding, provides evidence that *Safety Prediction* evokes dissociable circuits depending on whether the stimulus has self-relevance, and supports the hypothesis that the brain integrates threat and protective information to *Meta-represent* safety. The vmPFC emerged as a robust hub of human safety coding during safety estimation, including *Safety Prediction*, *Meta-representation*, *Recognition*, and *Value Updating*. The vmPFC showed specific tuning to protective information, supporting the importance of developing models of safety computing to expand beyond extinction of external threat.

During *Safety Prediction*, subjects were quicker to detect safety when presented with self-relevant protective stimuli compared to when presented with externally-relevant threat stimuli. Neurally, vmPFC activation parametrically increased as protection increased in safety value. Threat stimuli, in contrast, activated sensory and defensive neural systems. Despite equivalent objective probabilities for threat and protection, only protection evoked activation in the hypothesized vmPFC safety region. These findings point to the importance of self-relevant information in the human estimation of safety.

During *Safety Meta-representation*, subjects were again quicker to detect safety for protection. Neurally, subjects meta-represented the first stimulus when evaluating the second stimulus, despite the absence of perceptual information about the first stimulus. In response to threat stimuli, the same vmPFC region that activated to protection during *Safety Prediction* was activated. In response to protection stimuli, the same sensory regions of the visual cortex that activated to threat during *Safety Prediction* were activated. In other words, the pattern of activation at the second stimulus was inverted, which we interpret as meta-representation during safety integration. This interpretation is bolstered by our design: stimuli at the second stimulus pairing were perceptually identical and only differed in safety value as a function of their pair. Thus, the neural systems responding to increases in safety during *Meta-representation* were responding to changes in the safety value as a function of the first stimulus safety value and its resulting influence on the overall safety probability. Similar behavioral patterns emerged: subjects were more likely to update predictions during *Safety Meta-representation* when self-relevant protection conflicted with initial external threat information. On the other hand, when protection was presented first, subjects did not shift safety estimations. Our results fit with a broader body of literature showing that memory retrieval induces aspects of the pattern of neural activity evoked by the original stimulus presentation.^32,33^

Given the lack of information that classic univariate connectivity approaches have (i.e., PPI) we used informational connectivity to examine synchronization of voxels. Multivariate connectivity revealed a safety network consisting of the anterior and posterior vmPFC, dorsal and ventral striatum, caudodorsal ACC, and insula. The hippocampus, thalamus, and amygdala also emerged as connected to this core network, but with less consistency depending on stimulus relevance. In response to *Safety Prediction*, threat and protection networks showed a shift in hub organization from the posterior to anterior vmPFC. The dorsal striatum emerged as a core hub for *Safety Meta-representation*. The dorsal striatum has been previously linked to punishment-based avoidance, with dorsal striatum damage resulting in suboptimal defensive choice.^34^ We situate our findings with consideration of this prior work and interpret *Safety Meta-representation* as necessary to generate choices about defensive action. The ventral striatum and hippocampus were both connected to the safety network during protection evaluation but not during threat evaluation, suggesting a reward-like signal in response to self-relevant safety. This is consistent with our prior work demonstrating that safety conferred through protection is distinct from threat despite both occurring in aversive contexts.^35^ The PAG did not emerge as part of the safety network, consistent with its role in fast innate defensive reactions.^36,37^ It did however emerge preceding risk of shock both during outcomes for lost battles and under increasing danger during *Meta-representation*.

Battle outcomes were tested as *Safety Recognition* and served as the purest test of safety neural circuitry. Neural activation in response to safety certainty, when subjects won the battle and were 100% safe from shock compared to when they lost battles, increased in the vmPFC, striatum, and hippocampus. In response to lost battles when there was risk of electric shock, compared with won battles, subjects demonstrated increased engagement of canonical defensive circuitry in the PAG and bilateral insula. Activation of the striatum and hippocampus indicates subjects learned during outcomes but did not reengage these circuits during prediction. The vmPFC, however, was engaged in response to both *Safety Prediction and Recognition*. We interpret these differences to indicate a more general role of the vmPFC in recognizing and predicting safety, rather than tracking outcomes to reinforce learning.

The vmPFC was identified through searchlight analyses as involved in *Value Updating* for high-safety stimuli. The vmPFC was differentially engaged after learning about safety stimuli compared to naive viewing. Importantly, the stimuli in this study were all perceptually threatening and therefore could all have been interpreted as dangerous when subjects viewed them without knowledge of the Safety Estimation Task. Dangerous weapons have a general threat connotation and only take on a safety status when wielded to protect oneself. All animals presented as threats were shown to be attacking and angry (as opposed to a cuddly housecat). Thus, we expected that all stimuli would be represented as threatening before learning that the weapons provided safety to the subjects. Changes in multivariate vmPFC representation after experience with the stimuli provide converging evidence that the vmPFC integrates information about safety rather than processing a more general stimulus value.

Although this study makes a significant advance in understanding how the brain contributes to estimating safety, many questions remain. First, how universal is the role of the vmPFC in coding safety? This study used a model of predator-prey interactions. Although humans are not typically exposed to predation, everyday threats induce neurobiological and psychophysiological states like those observed under predatory threats.^3,13,38^ Using outwardly dangerous animals as threats eliminated interference from prior experience and allowed us to test the neural systems involved in *Safety Value Updating*. However, future work should examine safety coding during a diversity of threats including complex human interactions. Second, how do safety network communications evolve as a function of spatiotemporal dynamics? Our prior work shows that neural systems involved in defensive responding are dissociable along the threat imminence continuum.^36,39^ The task used in the current study was not designed to examine dynamic threat nor did it evoke escape behavior. Further work is needed to determine under what conditions vmPFC-supported cognition is unavailable. Lastly, our MRI sample size prevented the examination of brain-based individual differences in age and psychopathology. Adolescence is a critical time to study safety computations given the prevalence of anxiety disorders, changes in metacognitive abilities, poorer threat-safety discrimination compared with adults, and imbalance in amygdala–vmPFC contributions to safety processing.^27,40–43^ These features of adolescent development may result in impaired self-relevant safety processing.

This study demonstrates differential engagement of human neural circuitry as a function of safety relevance (external versus self), both when information was incomplete and during full information integration. The vmPFC coded protection during *Safety Recognition* and *Meta-representation*. Converging evidence of vmPFC involvement was observed during *Safety Recognition* and *Value Updating*. We identified a safety coding network that included subcortical and cortical regions involved in diverse processes including learning, reward valuation, and affect signaling. Our findings support assertions that the anterior vmPFC plays a role in integrating the value of self-relevant stimuli to influence the higher-order construction of affective processes, including safety.^2,44^

Beyond identifying how the human brain codes safety, our findings have potential implications for improving clinical interventions for anxiety. Current therapies focus on threat extinction but are ineffective for up to 50% of individuals.^45^ A major problem with studying safety through the lens of threat extinction is the assumption that safety is the inverse of threat (in the absence of the aversive event the stimulus itself becomes “safe”). This confounding association does not consider how safety fluctuates independent of threat, for example when protective resources can change safety while external threats remain unchanged. Current therapies target extinguishing fear responses to threats,^46^ but our data suggest focusing on self-relevant safety cues may be a promising therapeutic avenue. Also supporting a departure from extinction-focused approaches, recent work showed repetitive transcranial magnetic stimulation (rTMS) modulation of the anterior mPFC inhibited implicit fear reactions to learned threats.^47^ This is a departure from emphasis on the dorsolateral PFC as a regulatory hub, which may be limited to extinction paradigms.^48,49^ Intriguingly the mPFC is a hub of the brain’s Default Mode Network (DMN), which point to safety as an aspect of baseline human cognition. Psychopathologies, like anxiety, are often characterized by DMN dysfunction,^50,51^ which may mechanistically explain co-occurring deficits in safety estimation. Our findings provide a neuroscientifically-grounded framework of safety beyond threat extinction and set the stage for future research to better understand how the human brain adaptively codes safety.

## Methods

### Behavioral

One hundred thirteen human subjects completed the online version Safety Estimation Task. Subjects were recruited through Prolific, a recruitment and data collection platform that produces high-quality data.^52^ Seven subjects responded to fewer than 20% of trials and 6 subjects made safety choices that were inversely related to the safety continuum (i.e., judging safe stimuli as dangerous and dangerous stimuli as safe) resulting in an accuracy of >3 standard deviations (SD) below the group mean. Excluding these 13 subjects resulted in a final behavioral sample of 100 subjects (*M*age=29.20 years, *SD*=6.61, range=19-40, 50 females 51%).

### MRI

Thirty-one human subjects completed the Safety Estimation Task while undergoing functional MRI. Subjects were recruited through flyers and advertisements. One subject had +3SD below the group mean in accuracy and was excluded, resulting in a final MRI sample of 30 subjects (*M*age=27.83, *SD*=4.86, range 20-40 years, 15 females 50%,). One additional subject was excluded from analyses relating to the passive viewing task (*Safety Value Updating*) due to poor registration and dropout in the vmPFC, the primary area of interest. The full study was conducted for ∼90 minutes per day over 2 days.

### Inclusion and Exclusion Criteria

Inclusion criteria for both samples were age 18-40, fluent in English, and normal or corrected vision. The MRI sample was additionally required to have no psychiatric or neurological illness and be eligible for MRI, including having no metal contraindications.

### Ethics

All methodology was approved by the California Institute of Technology Internal Review Board, and all subjects consented to participation through a written consent form. Subjects were compensated for their time.

### Procedure

For MRI sessions, subjects first provided informed consent. Outside of the scanner, physiological equipment was attached and a shock workup procedure was conducted. During the shock workup procedure, shocks started at a low intensity and increased to the level the participant considered “uncomfortable but not painful” using a 0-10 discomfort scale (0 = “not at all,” 5 = “moderately,” and 10 = “very”; *M_session1_*= 4.87, *SD_session1_*= .34; *M_session2_*= 5.16, *SD_session2_*= .56). Shock intensity from session 1 was highly correlated with shock intensity from session 2 at *r*(31)=.93. Shocks were delivered using STMISOC with two LEAD110A (BIOPAC, Inc.) and two Telectrode T716 Ag/AgCl electrodes. The shock consisted of two pulses .03 sec apart delivered to the underside of the wrist 1-2 inches below the palm during outcome screens for lost battles.

While in the scanner (Fig. 1e), subjects first completed the passive viewing task to maintain ignorance to stimuli relevance. Next, subjects completed instructions for the Safety Estimation Task and 10 practice trials. During the first session, subjects completed a structural MPRAGE and 2 runs of the Safety Estimation Task. During the second session, subjects completed 2 runs of the Safety Estimation Task. After all Safety Estimation Task runs, subjects completed the passive viewing task again.

### Safety Estimation Task

During the Safety Estimation Task, subjects played a series of battles in which they attempted to defeat an animal with a weapon. Subjects did not receive a choice in animal or weapon and probabilities of winning or losing the battle were objectively determined (Fig. 1a-b). Stimuli were recognizable to reduce learning confounds (e.g., it is widely known that a grizzly is more dangerous than a goose). For each trial, subjects saw 2 images separated by a brief and variable interstimulus interval (ISI). Each pair contained one weapon and one animal, with presentation order counterbalanced. Subjects were told to indicate with a button press within 6 seconds while the stimulus was on screen whether they thought they would win or lose the battle against the animal with the weapon provided. Subjects were told “winning does not necessarily mean killing the other animal. You can interpret winning as defeating the other animal either because it retreats or because it is physically defeated”. Response to the first image presentation was based on partial information whereas response to the second image presentation was based on full information of the animal/weapon pair. Animals and weapons ranged in safety value on a 4-point continuum with matched contingencies (Fig. 1b). Likelihood of win/loss depended on the combined probability of the animal/weapon. For example, if subjects encountered a lion, they had an average 64.29% likelihood of losing the battle regardless of the weapon. That likelihood increased to 78.57% if subjects were equipped with a stick and reduced to 42.86% if subjects were equipped with a grenade. After both images were presented, subjects saw the outcome of the battle for 2 seconds. For the MRI sample, subjects had a 20% chance of receiving an electric shock to the wrist for every lost battle. Subjects were 100% safe from electric shock if they won the battle. For the behavioral sample, points were lost and gained depending on battle outcomes. Trials were separated by a variable inter-trial interval (ITI). All subjects completed 448 trials. The behavioral sample completed trials in a single session whereas the MRI sample completed 4 runs of 112 trials each run each over 2 days. The Safety Estimation Task was programmed using PsychoPy v2021.2.3.

### Stimuli Development

Prior to data collection, a series of stimulus development tests were conducted. Sixty subjects participated in stimuli development. Data from two subjects were excluded due to failure of attention checks resulting in a development sample of 58 (*M*age=23.07 years, *SD*=4.50, range=18-38, 39 females 67%). Twenty animals and 20 weapons were presented in paired head-to-head battles. For animal head-to-heads, subjects were asked to pick which animal they thought would win in a battle. Subjects were given the same instructions as the Safety Estimation Task: “winning does not necessarily mean killing the other animal. You can interpret winning as defeating the other animal either because it retreats or because it is physically defeated”. For weapon head-to-heads, subjects were asked to pick which object they thought was more powerful. Subjects were told, “You can think of this as choosing which weapon would win in a head-to-head battle”. The danger of each animal was also rated on a 0-100 scale (0=not at all dangerous, 100=extremely dangerous), and the power of each weapon was rated 0-100 (0=not at all powerful, 100=extremely powerful). Based on the results of these inquiries, a final set of 4 animals and 4 weapons were selected with two of each at the high end of the safety continuum and two of each at the low end. Stimuli were reduced to 4 based on scan time considerations and the number of trials needed for multivariate analyses. Lion and grizzly were rated as the most dangerous stimuli and cat and goose were rated as the second and third least dangerous stimuli (rat was selected as the least dangerous but ultimately excluded from the set to avoid conflating threat with disgust) (Supplemental Fig. S2a-b). The same rankings were reported for the head-to-head battles across all animals. The grenade and gun were rated as the most powerful weapons and as most likely to win head-to-head (Supplemental Fig. S2c-d). Fist and stick were rated in the bottom 30% of power ratings and bottom 20% of head-to-heads. Other weapons rated as less powerful were excluded due to concerns of unwieldy usage (i.e., rope) (Supplemental Fig. S2c-d).

### Safety Value Updating

Before and after the Safety Estimation Task, the MRI sample completed a passive viewing task. During the task, subjects saw animal and weapon images that were used in the Safety Estimation Task in a series of blocks. Subjects were instructed to look at each image carefully and that no decisions were required and no shocks would be administered. Images were presented for .5 seconds per image. Each image was followed by a .3 second ISI and each block was followed by a 12-second ITI. Images were presented 5x in each block. Six blocks were repeated 5x each. Blocks were comprised of high danger stimuli with high shock probability during the Safety Estimation Task, high safety stimuli with low shock probability during the Safety Estimation Task, high threat stimuli with high external threat value outside of the experimental environment, low threat stimuli that have low external threat value outside of the environmental experiment, low threat stimuli with low external threat value, weapons, and animals.

### Behavioral Models

Mixed effects models were fit using R (version 4.1.3) and the lme4 package (version 1.1.28).^53^ Frequentists probabilities were determined using the Satterthwaite method via lmerTest. General effect sizes are reported as 95% confidence intervals. Model effect sizes reported as *R*^2^ are conditional effects of variance explained by the entire model.^54^

### MRI Data Acquisition

Functional and structural data were acquired using a Siemens 3 Tesla Magnetom Prisma MRI scanner. For the acquisition of the functional images, we used a T2* weighted gradient EPI sequence. The repetition time (TR) was 1.12 s, the echo time (TE) was 30 milliseconds, the flip angle was 54 degrees, and the voxel resolution was 2 x 2 x 2 mm. A total of 512 slices were acquired in ascending interleaved order with a multiband acceleration factor of 4. Each functional run consisted of 1279 volumes.

For the structural data, we used a T1*-weighted MPRAGE sequence (image size 208 x 256 x 256 voxels, TR 2.55 s, TE .16 ms, flip angle 8, slice thickness=0.9 mm).

Stimuli were projected onto a flat screen mounted in the scanner bore. Participants viewed the screen using a mirror mounted on a 32-channel head coil. Extensive head padding was used to minimize participant head motion and to enhance comfort. Participants made their safety judgments with their right hand using a 4-finger-button response box.

### MRI Preprocessing

Raw data were converted from DICOM to BIDS format. Results included in this manuscript come from preprocessing performed using fMRIPrep 21.0.0^55,56^ (@fmriprep1; @fmriprep2; RRID:SCR_016216), which is based on Nipype 1.6.1 (@nipype1; @nipype2; RRID:SCR_002502). A B0-nonuniformity map (or fieldmap) was estimated based on two (or more) echo-planar imaging (EPI) references with ‘topup’ (@topup; FSL 6.0.5.1:57b01774). The T1-weighted (T1w) image was corrected for intensity non-uniformity (INU) with ‘N4BiasFieldCorrection’ [@n4], distributed with ANTs 2.3.3 [@ants, RRID:SCR_004757], and used as T1w-reference throughout the workflow. The T1w-reference was then skull-stripped with a Nipype implementation of the ‘antsBrainExtraction.sh’ workflow (from ANTs), using OASIS30ANTs as target template. Brain tissue segmentation of cerebrospinal fluid (CSF), white-matter (WM) and gray-matter (GM) was performed on the brain-extracted T1w using ‘fast’ [FSL 6.0.5.1:57b01774, RRID:SCR_002823, @fsl_fast]. Volume-based spatial normalization to one standard space (MNI152NLin2009cAsym) was performed through nonlinear registration with antsRegistration (ANTs 2.3.3), using brain-extracted versions of both T1w reference and the T1w template. The following template was selected for spatial normalization: ICBM 152 Nonlinear Asymmetrical template version 2009c [@mni152nlin2009casym, RRID:SCR_008796; TemplateFlow ID: MNI152NLin2009cAsym]. For each of BOLD run per subject, the following preprocessing was performed: First, a reference volume and its skull-stripped version were generated using a custom methodology of fMRIPrep. Head-motion parameters with respect to the BOLD reference (transformation matrices, and six corresponding rotation and translation parameters) are estimated before any spatiotemporal filtering using ‘mcflirt’ [FSL 6.0.5.1:57b01774, @mcflirt]. BOLD runs were slice-time corrected to 0.52s (0.5 of slice acquisition range 0s-1.04s) using ‘3dTshift’ from AFNI [@afni, RRID:SCR_005927]. The BOLD time-series (including slice-timing correction when applied) were resampled onto their original, native space by applying the transforms to correct for head-motion. These resampled BOLD time-series will be referred to as ‘preprocessed BOLD’. The BOLD reference was then co-registered to the T1w reference using ‘mri_coreg’ (FreeSurfer) followed by ‘flirt’ [FSL 6.0.5.1:57b01774, @flirt] with the boundary-based registration [@bbr] cost-function.

Co-registration was configured with six degrees of freedom. Several confounding time-series were calculated based on the preprocessed BOLD: framewise displacement (FD), DVARS, and three region-wise global signals. FD was computed using two formulations following Power (absolute sum of relative motions, @power_fd_dvars) and Jenkinson (relative root mean square displacement between affines, @mcflirt). FD and DVARS are calculated for each functional run, both using their implementations in Nipype [following the definitions by @power_fd_dvars]. The three global signals are extracted within the CSF, the WM, and the whole-brain masks. Additionally, a set of physiological regressors were extracted to allow for component-based noise correction [CompCor, @compcor]. Principal components are estimated after high-pass filtering the preprocessed BOLD time-series (using a discrete cosine filter with 128s cut-off) for the two CompCor variants: temporal (tCompCor) and anatomical (aCompCor). tCompCor components are then calculated from the top 2% variable voxels within the brain mask. For aCompCor, three probabilistic masks (CSF, WM and combined CSF+WM) are generated in anatomical space. aCompCor masks are subtracted a mask of pixels that likely contain a volume fraction of GM. This mask is obtained by thresholding the corresponding partial volume map at 0.05, and it ensures components are not extracted from voxels containing a minimal fraction of GM. Finally, these masks are resampled into BOLD space and binarized by thresholding at 0.99 (as in the original implementation). Components are also calculated separately within the WM and CSF masks. For each CompCor decomposition, the k components with the largest singular values are retained, such that the retained components’ time series are sufficient to explain 50 percent of variance across the nuisance mask (CSF, WM, combined, or temporal). The remaining components are dropped from consideration. The head-motion estimates calculated in the correction step were also placed within the corresponding confounds file. The confound time series derived from head motion estimates and global signals were expanded with the inclusion of temporal derivatives and quadratic terms for each [@confounds_satterthwaite_2013]. Frames that exceeded a threshold of 0.5 mm FD or 1.5 standardised DVARS were annotated as motion outliers. The BOLD time-series were resampled into standard space, generating a preprocessed BOLD run in MNI152NLin2009cAsym space. First, a reference volume and its skull-stripped version were generated using a custom methodology of fMRIPrep. All resamplings can be performed with a single interpolation step by composing all the pertinent transformations (i.e. head-motion transform matrices, susceptibility distortion correction when available, and co-registrations to anatomical and output spaces). Gridded (volumetric) resamplings were performed using ‘antsApplyTransforms’ (ANTs), configured with Lanczos interpolation to minimize the smoothing effects of other kernels [@lanczos]. Non-gridded (surface) resamplings were performed using ‘mri_vol2surf’ (FreeSurfer). Many internal operations of fMRIPrep use Nilearn 0.8.1 [@nilearn, RRID:SCR_001362], mostly within the functional processing workflow. For more details of the pipeline, see https://fmriprep.readthedocs.io/en/latest/workflows.html.

### Univariate Analysis

For univariate group-level fMRI analyses, FSL Randomise with 5000 permutations was used. Randomise uses a permutation-based statistical inference that does not rely on a Gaussian distribution (Nichols and Holmes, 2002). A statistical threshold of *p* < .05, corrected for multiple comparisons with familywise error correction (FWE) and threshold-free cluster enhancement (TFCE), was used for analyses. TFCE helps identify significant clusters without defining an initial cluster-forming threshold or carrying out a large amount of data smoothing (Smith and Nichols, 2009). Conjunction analyses were conducted using the easythresh_conj script in FSL^57^ and recommended thresholds (*Z* > 2.3, cluster size *p* < .05) to identify regions commonly activated for *Safety Prediction* and *Meta-representation*. MRIcron was used for visualization.

### Informational Connectivity Analysis

IC analyses were conducted using the IC Toolbox in Matlab.^58^ An advantage of IC over univariate functional connectivity is that IC utilizes all patterns of responses within regions to code information that is lost by averaging, which identifies functional connections that cannot be found in univariate functional connectivity analyses^23,24^. Furthermore, IC allows us to test regional interactions in terms of specific experimental conditions^24^ such as estimating safety in response to threat (animals) versus protection (weapons). IC was measured between every pair of the ten ROIs: hippocampus, thalamus, amygdala, PAG, anterior insula, caudodorsal ACC, dorsal striatum, ventral striatum, posterior vmPFC, anterior vmPFC (Fig. 4a). Network-based statistics were calculated for multi-voxel pattern synchronization changes as a function of *Safety Prediction* during first stimulus presentation to partial information (Fig. 4b) and *Safety Meta-representation* during second stimulus presentation as a function of paired stimuli (Fig. 4c). To identify hubs connecting regions within the networks identified, we computed the betweenness centralities of each region, which represents the fraction of all shortest paths that contain a specific node (Fig. 4b-c). BCs of 0 are not plotted.

### Regions of Interest

An independent vmPFC mask was defined and used for the *Safety Value Updating* searchlight analysis. The vmPFC mask was obtained via neurovault (https://identifiers.org/neurovault.image:132836) consisting of 152 subjects and 4233 voxels. The mask was transformed to standard space (MNI152NLin2009cAsym) using ‘flirt’ [FSL 6.0.5.1:57b01774, @flirt]. For IC analyses, ROIs were defined based on the Harvard-Oxford cortical and subcortical structural atlases, except for the PAG which was defined using Neurosynth meta-analysis (https://neurosynth.org/; “periaqueductal” with an association test).

### Preregistration

Hypotheses and methods were preregistered on the Open Science Framework (OSF), https://osf.io/hw3r9.

## Data availability

Task code and raw data are available through OSF, https://osf.io/hw3r9.

## Acknowledgments

During the study, DM was supported by a US National Institute of Mental Health (NIMH) grant NIMH R01MH133730, Templeton Foundation grant TWCF0366, and SMT was supported by a National Science Foundation SBE Postdoctoral Research Fellowship 2203522. SMT is currently supported by a Brain & Behavior Research Foundation Young Investigator Grant 30788.

## Contributions

Design of the study: SMT, DM. Behavioral data collection: SMT. Neuroimaging data collection: SMT, WD. Data processing, quality control, management: SMT. Behavioral data analysis: SMT, WD. Neuroimaging data analysis: SMT, JC, with the input of BZ. Data interpretation and generation of figures: SMT. Writing original draft: SMT. Reviewing, editing: SMT, DM. Approval: All authors.

## Competing Interests

The authors declare no competing interests.

---

**Fig S1.**
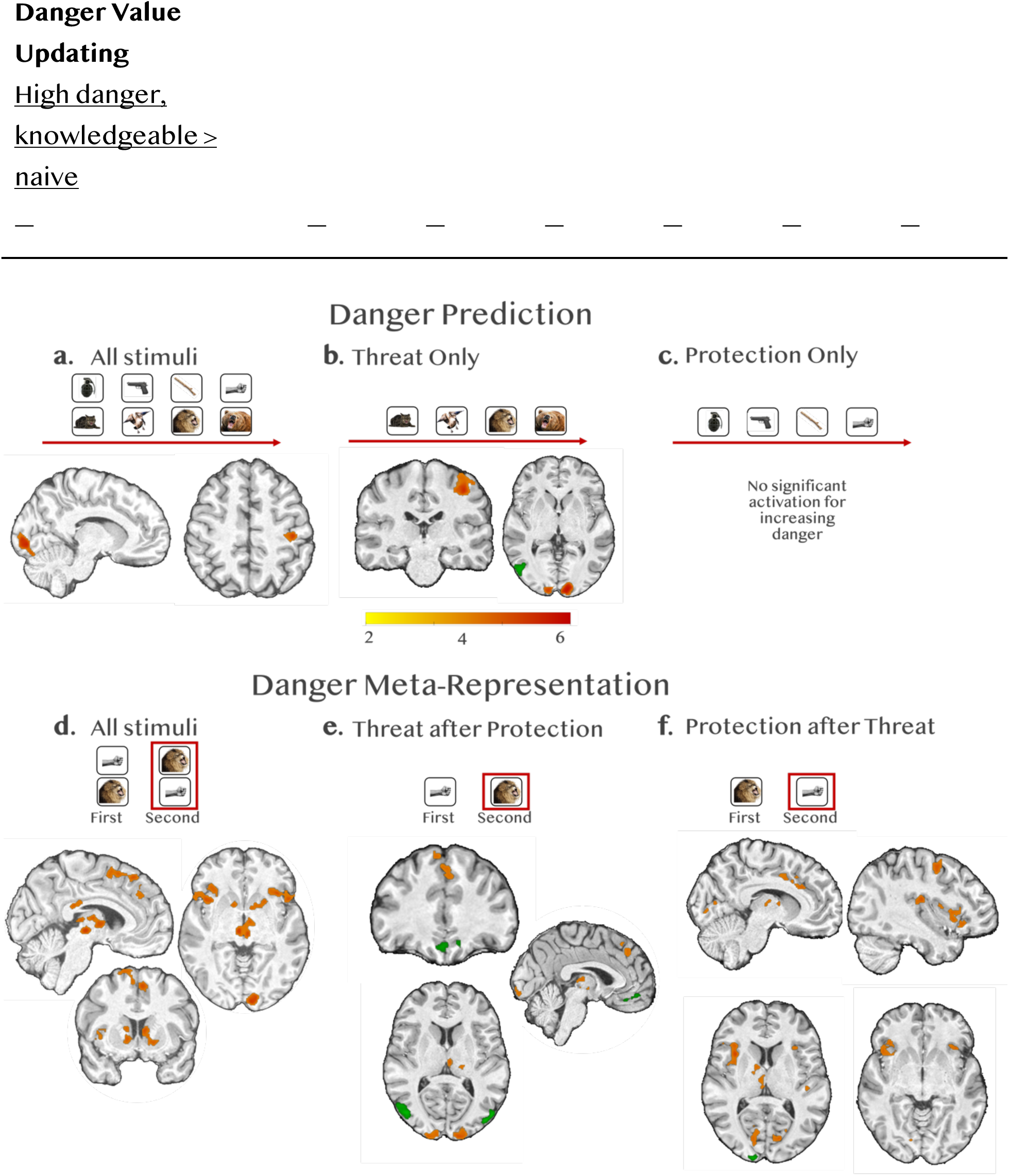
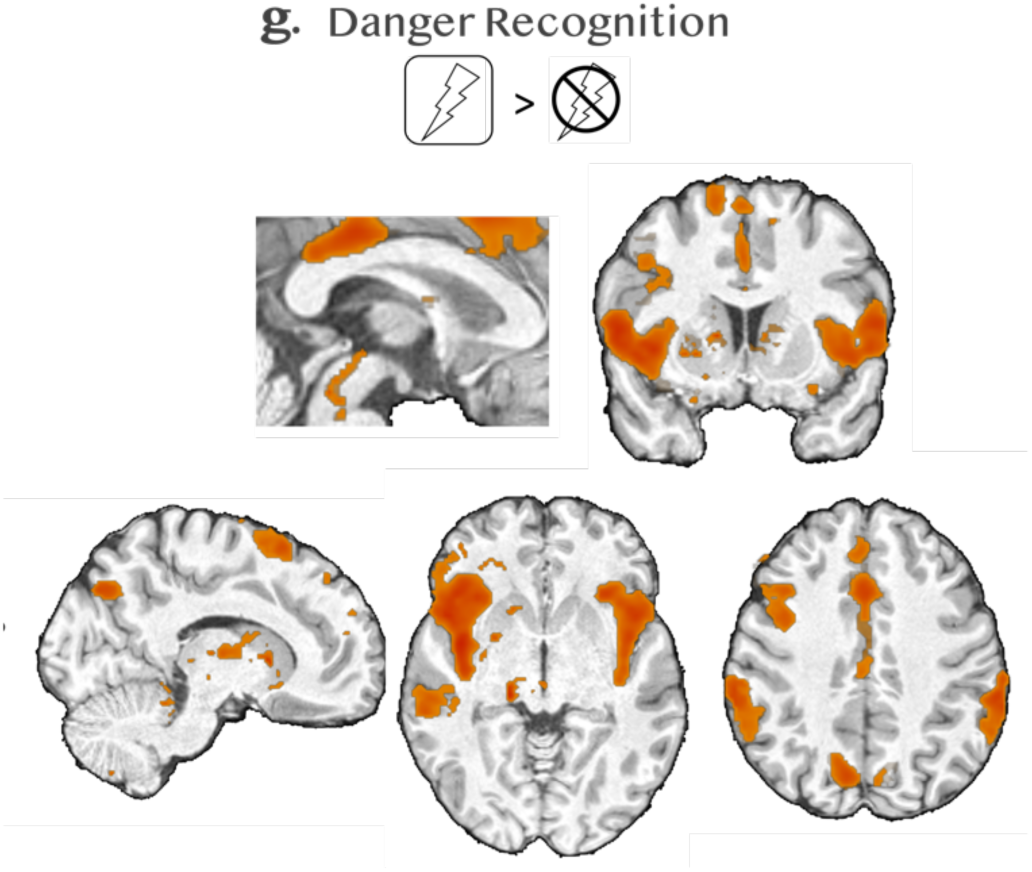
Visualization of Univariate Results. Parametric increases in whole-brain neural activity that track decreases in objective safety value (increases in danger) of stimuli. All analyses in Figure 3 were conducted using FSL Randomise, TFCE, FWE-corrected *p*<.05. Color bar indicates t-intensity values. **Danger Prediction:** First stimulus presentation during the Safety Estimation Task. Activation increased as danger increased based on average shock probability of each stimulus. **(a)** Threat and Protection collapsed, **(b)** Threat only, **(c)** Protection only. **Danger Meta-representation:** Second stimulus presentation during the Safety Estimation Task. Danger increases were based on comparison with the average shock value of the stimulus. **(d)** Threat and Protection collapsed, **(e)** Threat only, **(f**) Protection only. **Danger Recognition: (g)** Neural activation in response to unsuccessful battles during the outcome screen that indicated potential for electric shock (20%) compared to certain safety from shock (100%). All analyses were conducted using FSL Randomise, TFCE, FWE-corrected *p*<.05. Color bar indicates t-intensity values.

**Fig S2.**
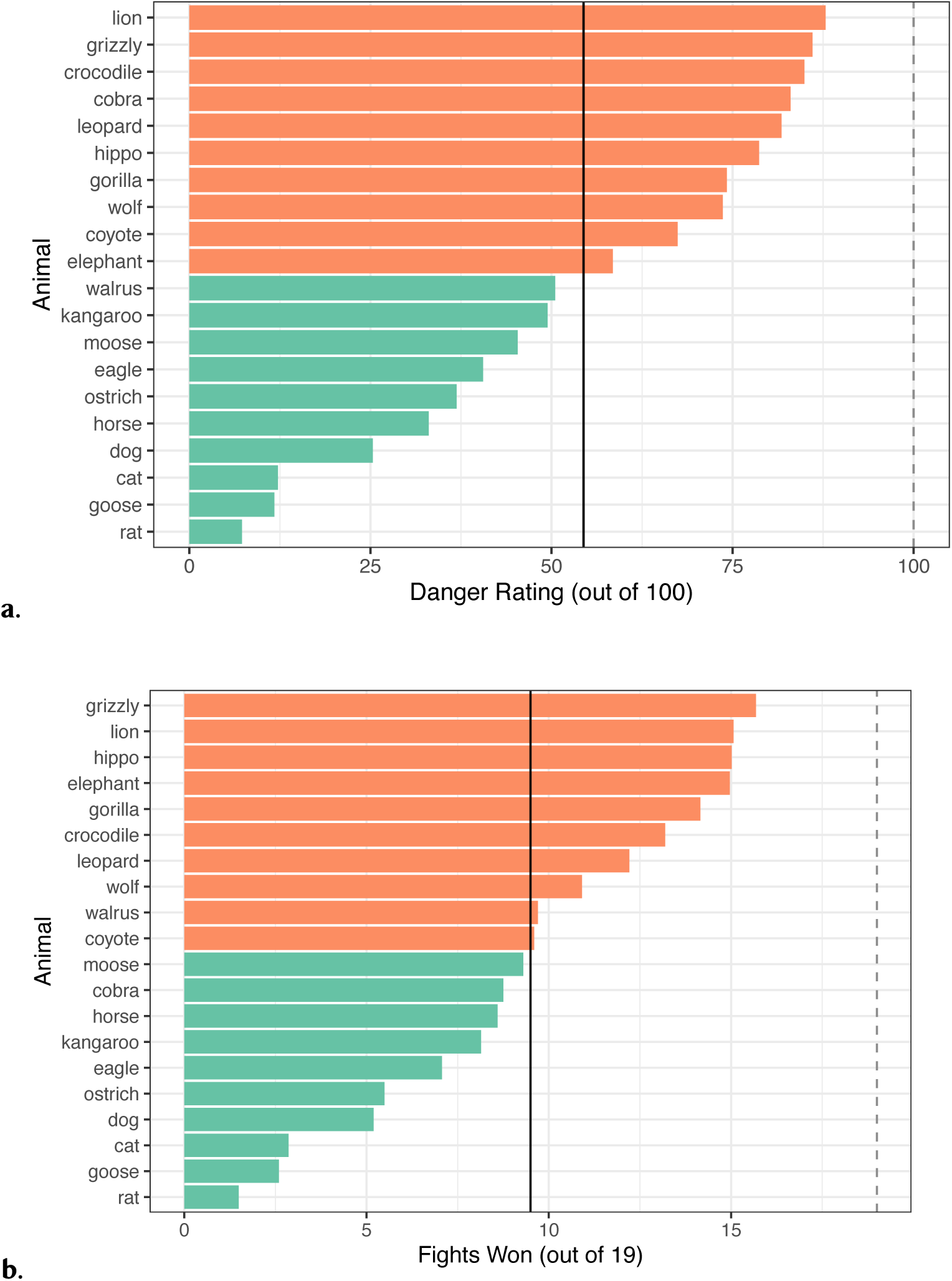
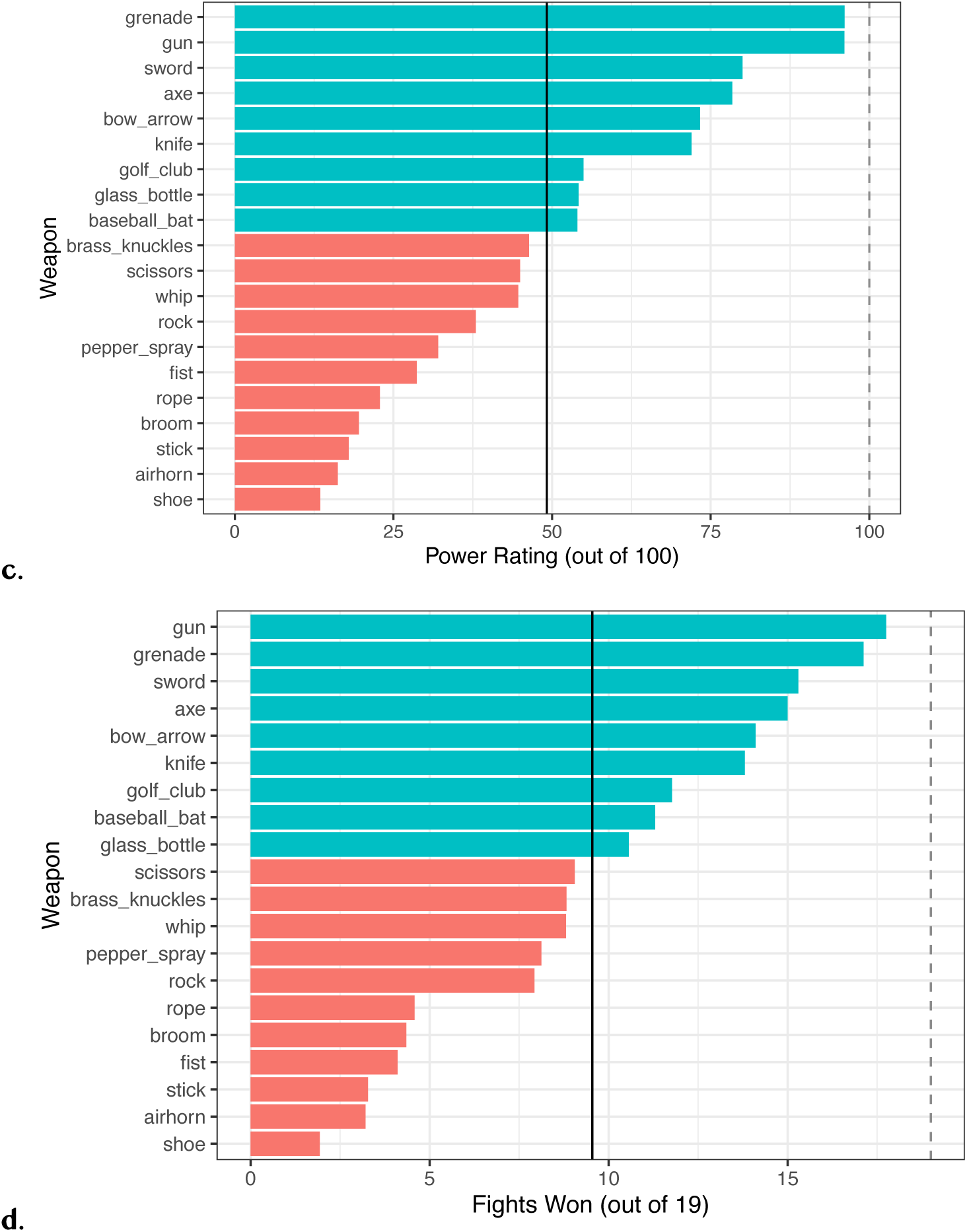
Results of stimuli development questions related to level of danger (animals) and power (weapons) as well as when items were pitted in head-to-head battles with all other stimuli of the same type.

**Table S1.**
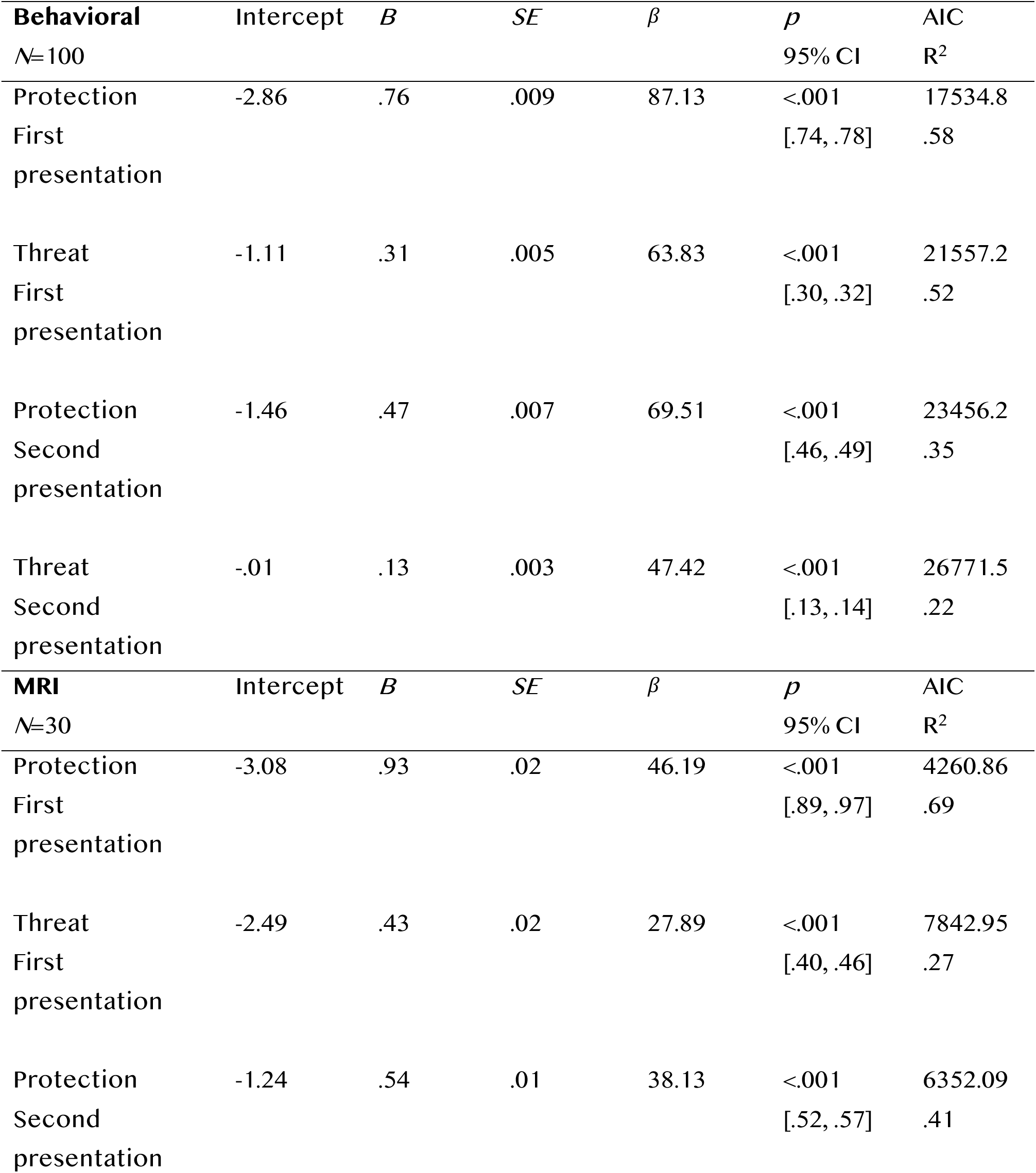
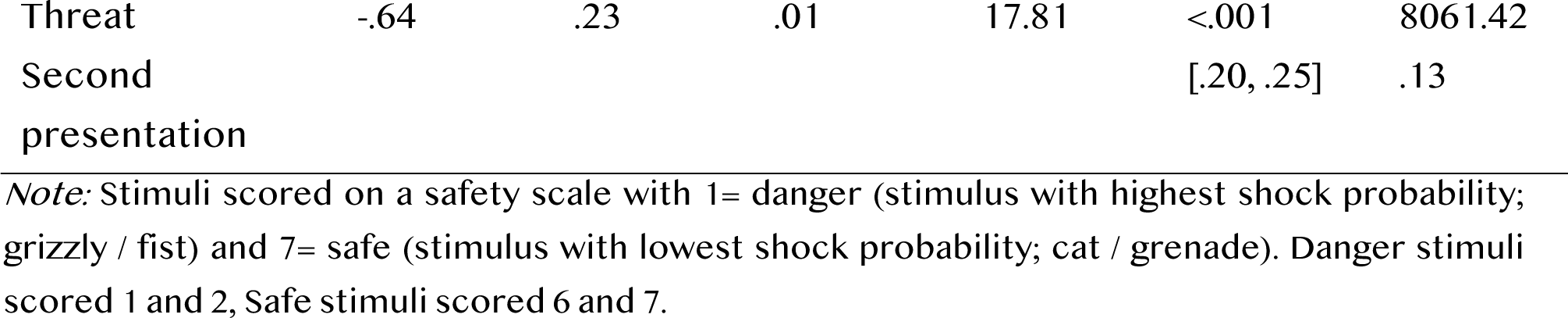
Mixed effects models predicting safety prediction (probability of choosing ‘win’) from stimuli safety value, split by stimulus order and type.

**Table S2.**
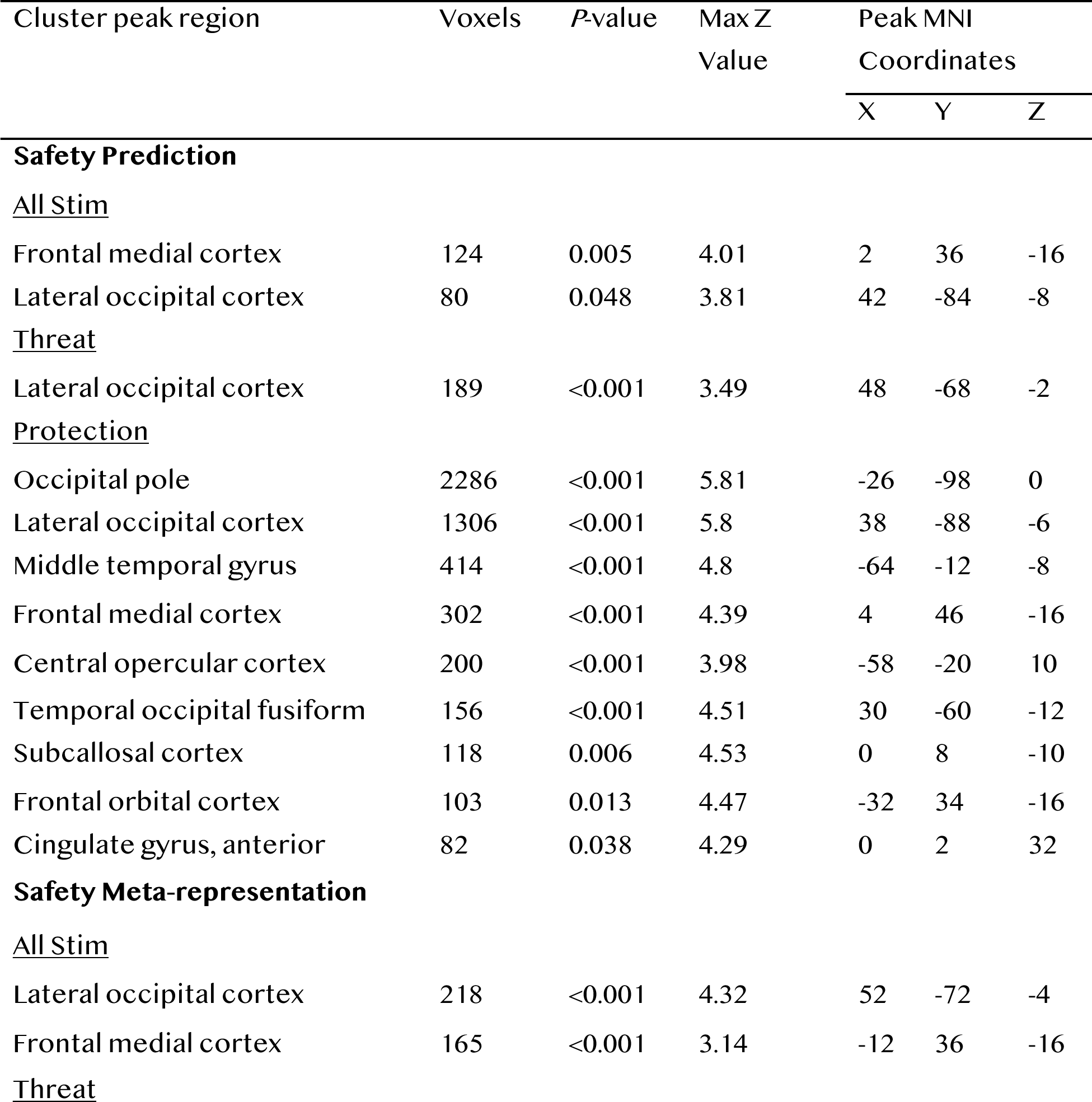
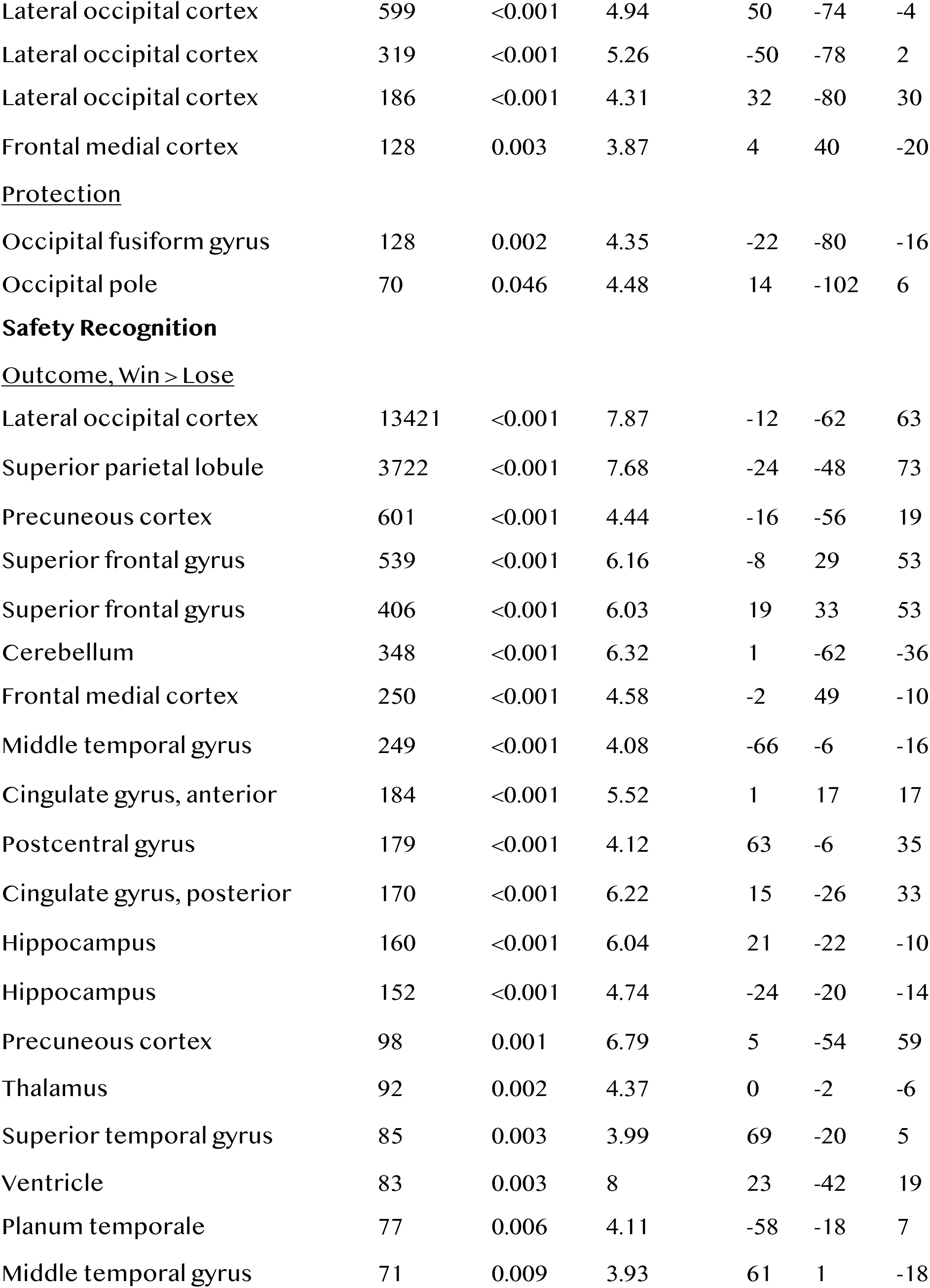
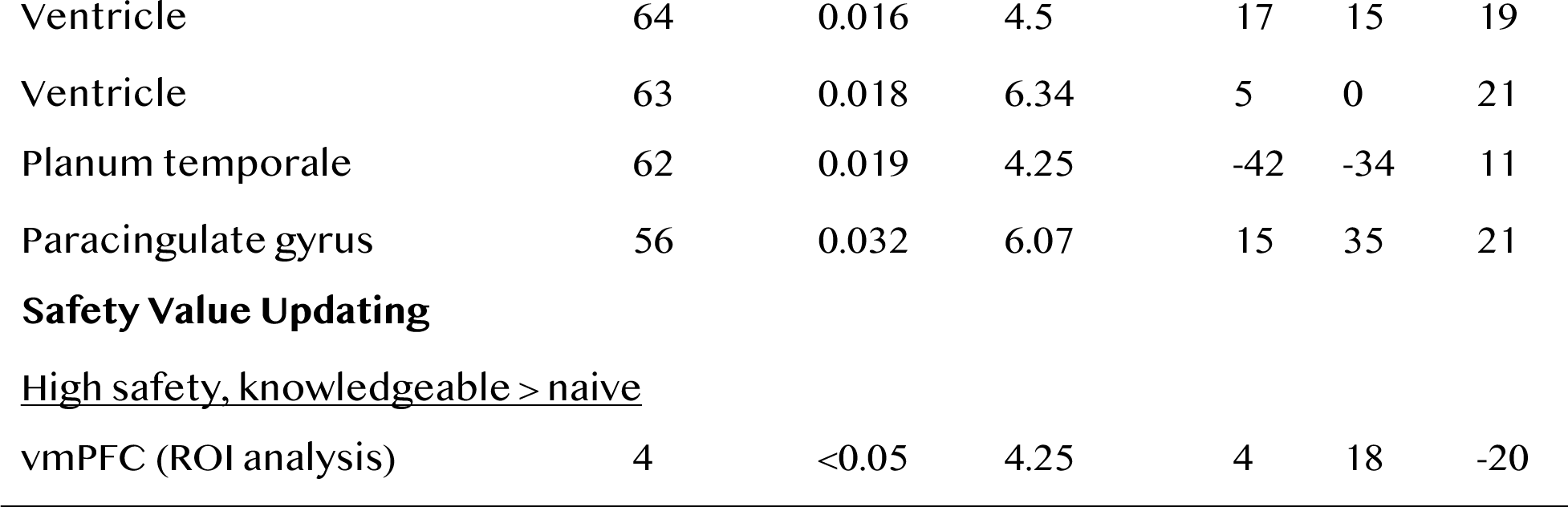
Neural response to safety. Significant clusters from group level whole-brain univariate analyses.

**Table S3.**
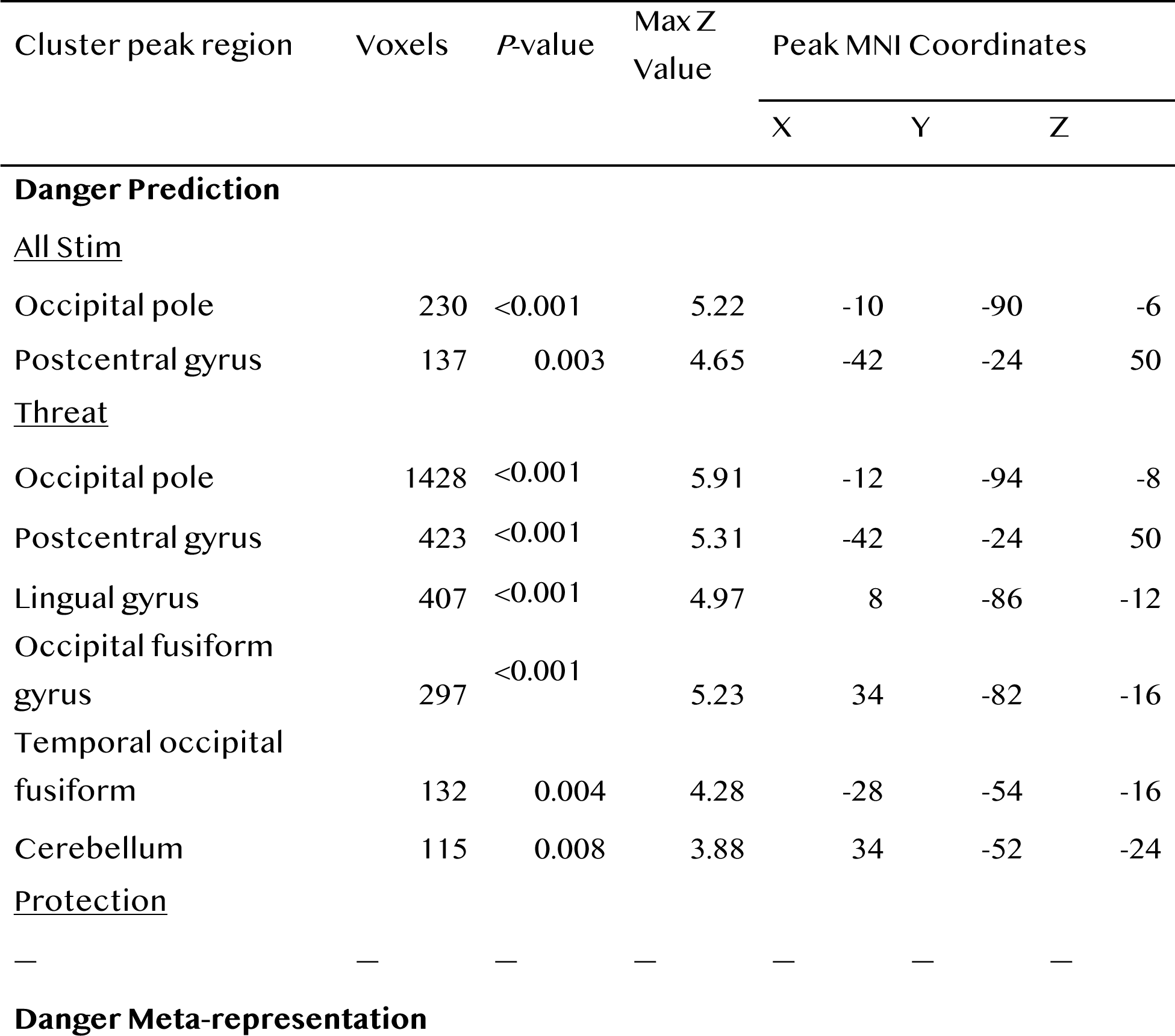
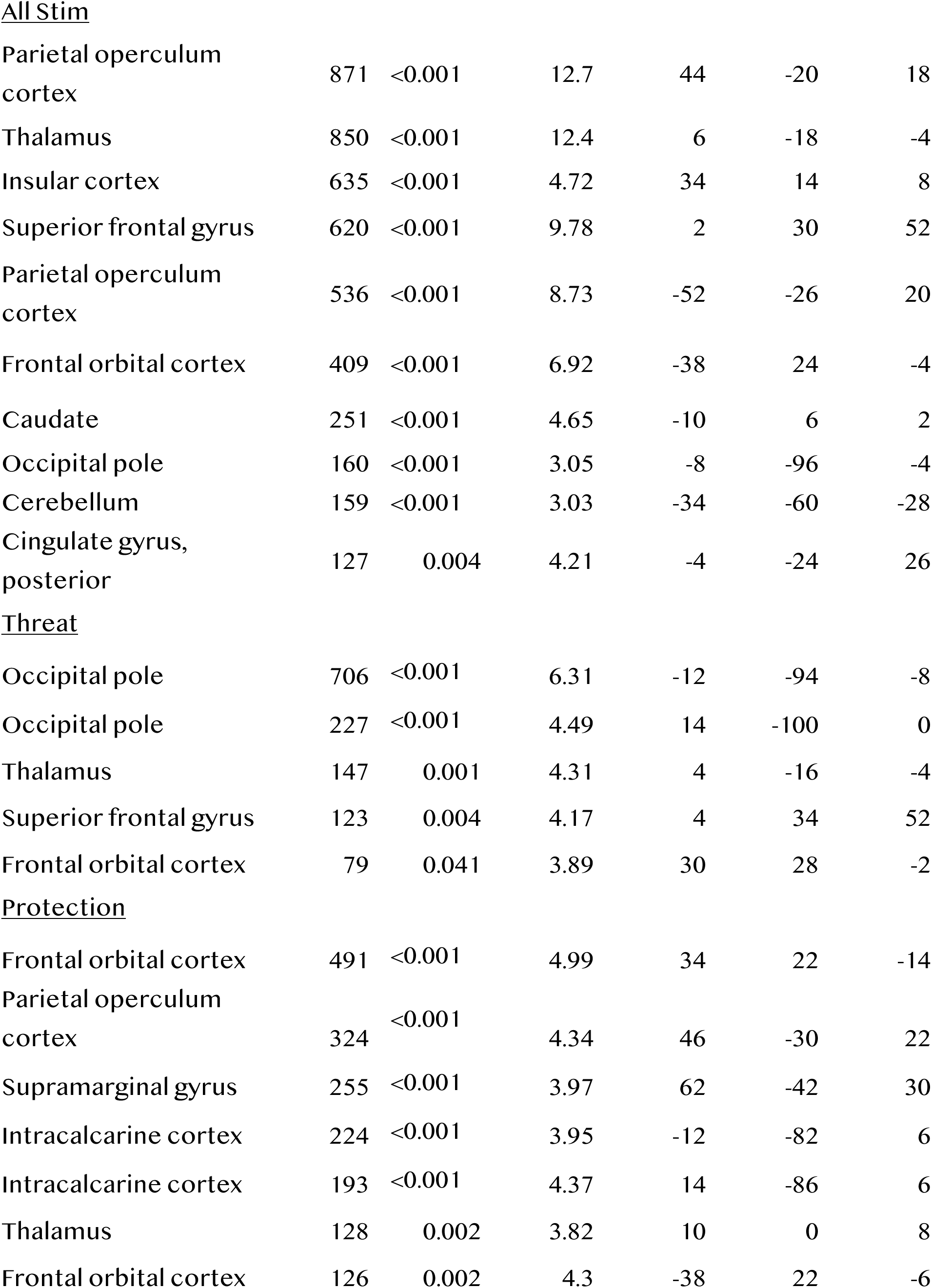
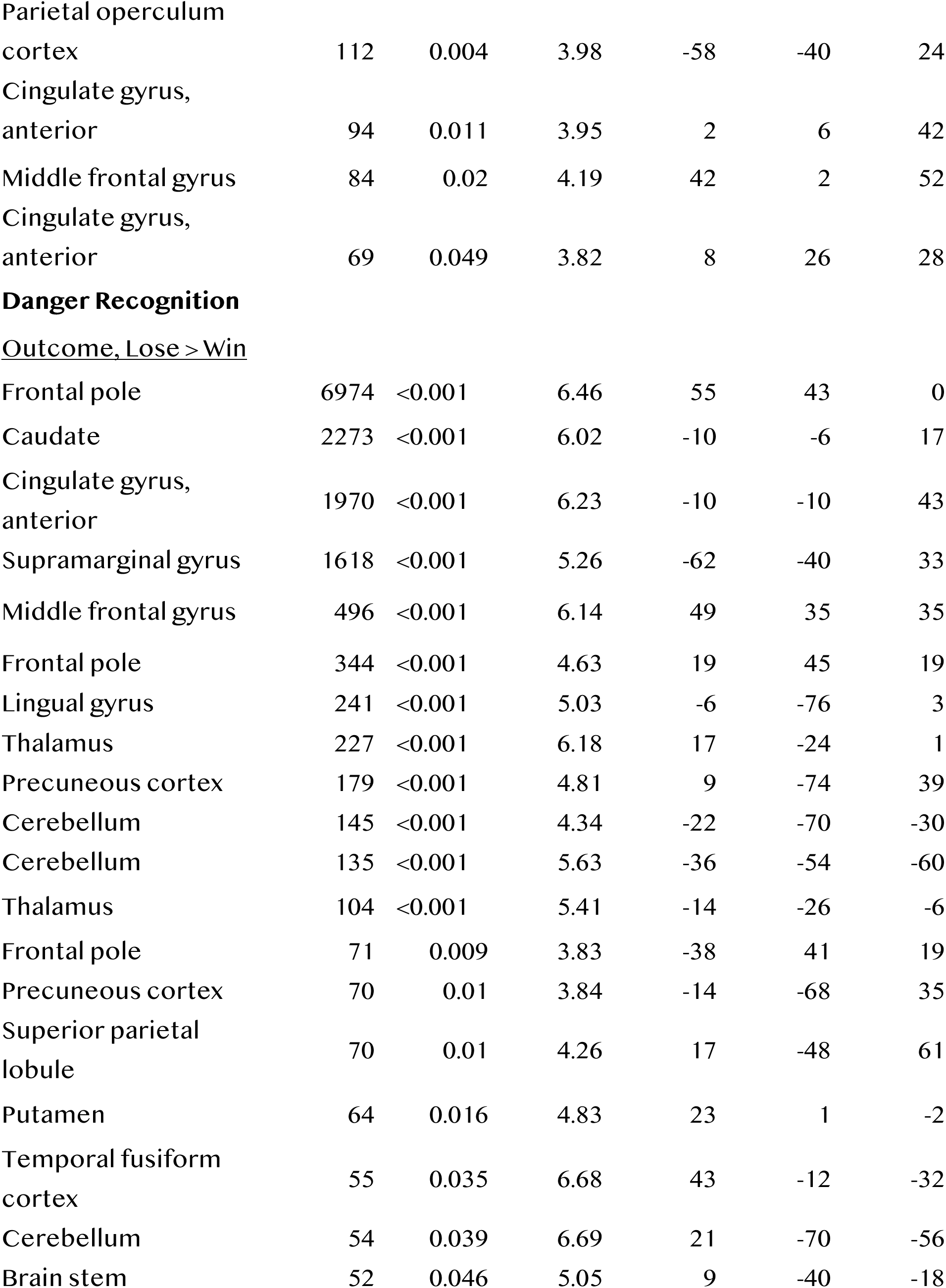
Neural response to danger. Significant clusters from group level whole-brain univariate analyses.

## Notes

### Competing Interest Statement

The authors have declared no competing interest.

## Reference

1. Neuberg SL, Kenrick DT, Schaller M. Human Threat Management Systems: Self-Protection and Disease Avoidance. Neuroscience and biobehavioral reviews. 2011;35(4):1042.

2. Tashjian SM, Zbozinek TD, Mobbs D. A Decision Architecture for Safety Computations. Trends in Cognitive Sciences. 2021;25(5):342–354.

3. Mobbs D, Trimmer PC, Blumstein DT, Dayan P. Foraging for foundations in decision neuroscience: insights from ethology. Nature reviews. Neuroscience. 2018;19(7):419–427.

4. Lima SL, Dill LM. Behavioral decisions made under the risk of predation: a review and prospectus. Canadian Journal of Zoology. 1990;68(4):619–640.

5. Kveraga K, Ghuman AS, Bar M. Top-down predictions in the cognitive brain. Brain and cognition. 2007;65(2):145.

6. Rao RPN, Ballard DH. Predictive coding in the visual cortex: a functional interpretation of some extra-classical receptive-field effects. Nature neuroscience. 1999;2(1):79–87.

7. De-Wit L, Machilsen B, Putzeys T. Predictive Coding and the Neural Response to Predictable Stimuli. The Journal of Neuroscience. 2010;30(26):8702.

8. Harrison LA, Ahn C, Adolphs R. Exploring the Structure of Human Defensive Responses from Judgments of Threat Scenarios. PloS one. 2015;10(8).

9. Headley DB, Kanta V, Kyriazi P, Paré D. Embracing Complexity in Defensive Networks. Neuron. 2019;103(2):189–201.

10. Tovote P, Esposito MS, Botta P, Chaudun F, Fadok JP, Markovic M, Wolff SBE, Ramakrishnan C, Fenno L, Deisseroth K, et al. Midbrain circuits for defensive behaviour. Nature. 2016;534(7606):206–212.

11. Halladay LR, Blair HT. Distinct ensembles of medial prefrontal cortex neurons are activated by threatening stimuli that elicit excitation vs. inhibition of movement. Journal of Neurophysiology. 2015;114(2):793–807.

12. Moneta N, Garvert MM, Heekeren HR, Schuck NW. Task state representations in vmPFC mediate relevant and irrelevant value signals and their behavioral influence. Nature Communications. 2023;14(1):1–21.

13. Mobbs D, Headley DB, Ding W, Dayan P. Space, Time, and Fear: Survival Computations along Defensive Circuits. Trends in Cognitive Sciences. 2020;24(3):228–241.

14. Chalk M, Marre O, Tkačik G. Toward a unified theory of efficient, predictive, and sparse coding. Proceedings of the National Academy of Sciences of the United States of America. 2018;115(1):186–191.

15. Eldar E, Lièvre G, Dayan P, Dolan RJ. The roles of online and offline replay in planning. eLife. 2020;9:1–23.

16. Benoit RG, Szpunar KK, Schacter DL. Ventromedial prefrontal cortex supports affective future simulation by integrating distributed knowledge. Proceedings of the National Academy of Sciences of the United States of America. 2014;111(46):16550–16555.

17. Battaglia S, Garofalo S, di Pellegrino G, Starita F. Revaluing the role of vmPFC in the acquisition of pavlovian threat conditioning in humans. Journal of Neuroscience. 2020;40(44):8491–8500.

18. Laing PAF, Vervliet B, Dunsmoor JE, Harrison BJ. Pavlovian safety learning: an integrative theoretical review. [PREPRINT] https://osf.io/preprints/psyarxiv/p8xnw

19. Harrison BJ, Fullana MA, Via E, Soriano-Mas C, Vervliet B, Martínez-Zalacaín I, Pujol J, Davey CG, Kircher T, Straube B, et al. Human ventromedial prefrontal cortex and the positive affective processing of safety signals. Neuroimage. 2017;152:12–18.

20. Savage HS, Davey CG, Fullana MA, Harrison BJ. Clarifying the neural substrates of threat and safety reversal learning in humans. NeuroImage. 2020;207.

21. Battaglia S. Neurobiological advances of learned fear in humans. Advances in clinical and experimental medicine : official organ Wroclaw Medical University. 2022;31(3).

22. Battaglia S, Harrison BJ, Fullana MA. Does the human ventromedial prefrontal cortex support fear learning, fear extinction or both? A commentary on subregional contributions. Molecular Psychiatry. 2021;27(2):784–786.

23. Kveraga K, Ghuman AS, Bar M. Top-down predictions in the cognitive brain. Brain and cognition. 2007;65(2):145.

24. Frewen P, Schroeter ML, Riva G, Cipresso P, Fairfield B, Padulo C, Kemp AH, Palaniyappan L, Owolabi M, Kusi-Mensah K, et al. Neuroimaging the consciousness of self: Review, and conceptual-methodological framework. Neuroscience and biobehavioral reviews. 2020;112:164–212.

25. Hartley CA, Gorun A, Reddan MC, Ramirez F, Phelps EA. Stressor controllability modulates fear extinction in humans. Neurobiology of learning and memory. 2014;113:149–156.

26. Sangha S, Diehl MM, Bergstrom HC, Drew MR. Know safety, no fear. Neuroscience and Biobehavioral Reviews. 2020;108:218–230.

27. Odriozola P, Phil M, Gee DG. Learning About Safety: Conditioned Inhibition as a Novel Approach to Fear Reduction Targeting the Developing Brain. American Journal of Psychiatry. 2021;178:136–155.

28. Gilbert P. Threat, safety, safeness and social safeness 30 years on: Fundamental dimensions and distinctions for mental health and well-being. The British journal of clinical psychology. 2024.

29. Apergis-Schoute AM, Gillan CM, Fineberg NA, Fernandez-Egea E, Sahakian BJ, Robbins TW. Neural basis of impaired safety signaling in Obsessive Compulsive Disorder. Proceedings of the National Academy of Sciences of the United States of America. 2017;114(12):3216–3221.

30. Laing PAF, Steward T, Davey CG, Felmingham KL, Fullana MA, Vervliet B, Greaves MD, Moffat B, Glarin RK, Harrison BJ. Cortico-Striatal Activity Characterizes Human Safety Learning via Pavlovian Conditioned Inhibition. Journal of Neuroscience. 2022;42(25):5047– 5057.

31. Gonzalez ST, Fanselow MS. The role of the ventromedial prefrontal cortex and context in regulating fear learning and extinction. Psychology & Neuroscience. 2020;13(3):459.

32. Wimmer GE, Shohamy D. Preference by association: how memory mechanisms in the hippocampus bias decisions. Science. 2012:270–273.

33. Kurth-Nelson Z, Barnes G, Sejdinovic D, Dolan R, Dayan P. Temporal structure in associative retrieval. eLife. 2015;4(4):4919.

34. Palminteri S, Justo D, Jauffret C, Pavlicek B, Dauta A, Delmaire C, Czernecki V, Karachi C, Capelle L, Durr A, et al. Critical roles for anterior insula and dorsal striatum in punishment-based avoidance learning. Neuron. 2012;76(5):998–1009.

35. Tashjian SM, Wise T, Mobbs D. Model-based prioritization for acquiring protection. PLOS Computational Biology. 2022;18(12):e1010805.

36. Qi S, Hassabis D, Sun J, Guo F, Daw N, Mobbs D. How cognitive and reactive fear circuits optimize escape decisions in humans. Proceedings of the National Academy of Sciences of the United States of America. 2018;115(12):3186–3191.

37. Mobbs D, Headley DB, Ding W, Dayan P. Space, Time, and Fear: Survival Computations along Defensive Circuits. Trends in Cognitive Sciences. 2020;24(3):228–241.

38. Mobbs D, Hagan CC, Dalgleish T, Silston B, Prévost C. The ecology of human fear: Survival optimization and the nervous system. Frontiers in Neuroscience. 2015;9:121062.

39. Qi S, Cross L, Wise T, Sui X, O’doherty J, Mobbs D. The Role of the Medial Prefrontal Cortex in Spatial Margin of Safety Calculations. Journal of Neuroscience. (in press).

40. Lau JY, Britton JC, Nelson EE, Angold A, Ernst M, Goldwin M, Grillon C, Leibenluft E, Lissek S, Norcross M, et al. Distinct neural signatures of threat learning in adolescents and adults. Proceedings of the National Academy of Sciences of the United States of America. 2011;108(11):4500–4505.

41. Gee DG, Humphreys KL, Flannery J, Goff B, Telzer EH, Shapiro M, Hare TA, Bookheimer SY, Tottenham N. A developmental shift from positive to negative connectivity in human amygdala-prefrontal circuitry. The Journal of Neuroscience. 2013;33(10):4584–4593.

42. Paus T, Keshavan M, Giedd JN. Why do many psychiatric disorders emerge during adolescence? Nature Reviews Neuroscience. 2008;9(12):947–957.

43. Wilbrecht L, Davidow JY. Goal-directed learning in adolescence: neurocognitive development and contextual influences. Nature Reviews Neuroscience. 2024;25(3):176–194.

44. Roy M, Shohamy D, Wager TD. Ventromedial prefrontal-subcortical systems and the generation of affective meaning. Trends in Cognitive Sciences. 2012;16(3):147–156.

45. Springer KS, Levy HC, Tolin DF. Remission in CBT for adult anxiety disorders: A meta-analysis. Clinical psychology review. 2018;61:1–8.

46. Craske MG, Sandman CF, Stein MB. How can neurobiology of fear extinction inform treatment? Neuroscience & Biobehavioral Reviews. 2022;143:104923.

47. Manassero E, Concina G, Caraig MCC, Sarasso P, Salatino A, Ricci R, Sacchetti B. Medial anterior prefrontal cortex stimulation downregulates implicit reactions to threats and prevents the return of fear. eLife. 2024;13.

48. Dunsmoor JE, Bandettini PA, Knight DC. Neural correlates of unconditioned response diminution during Pavlovian conditioning. NeuroImage. 2008;40(2):811.

49. Marković V, Vicario CM, Yavari F, Salehinejad MA, Nitsche MA. A Systematic Review on the Effect of Transcranial Direct Current and Magnetic Stimulation on Fear Memory and Extinction. Frontiers in Human Neuroscience. 2021;15:655947.

50. Wen Z, Seo J, Pace-Schott EF, Milad MR. Abnormal dynamic functional connectivity during fear extinction learning in PTSD and anxiety disorders. Molecular Psychiatry. 2022;27(4):2216–2224.

51. Coutinho JF, Fernandesl SV, Soares JM, Maia L, Gonçalves ÓF, Sampaio A. Default mode network dissociation in depressive and anxiety states. Brain Imaging and Behavior. 2016;10(1):147–157.

52. Peer E, Brandimarte L, Samat S, Acquisti A. Beyond the Turk: Alternative platforms for crowdsourcing behavioral research. Journal of Experimental Social Psychology. 2017;70:153–163.

53. Bates D, Mächler M, Bolker BM, Walker SC. Fitting Linear Mixed-Effects Models Using lme4. Journal of Statistical Software. 2015;67(1):1–48.

54. Nakagawa S, Johnson PCD, Schielzeth H. The coefficient of determination R2 and intra-class correlation coefficient from generalized linear mixed-effects models revisited and expanded. Journal of The Royal Society Interface. 2017;14(134).

55. Esteban O, Markiewicz CJ, Blair RW, Moodie CA, Isik AI, Erramuzpe A, Kent JD, Goncalves M, DuPre E, Snyder M, et al. fMRIPrep: a robust preprocessing pipeline for functional MRI. Nature Methods. 2019;16(1):111–116.

56. Esteban O, Ciric R, Finc K, Blair RW, Markiewicz CJ, Moodie CA, Kent JD, Goncalves M, DuPre E, Gomez DEP, et al. Analysis of task-based functional MRI data preprocessed with fMRIPrep. Nature Protocols. 2020;15(7):2186–2202.

57. Nichols T, Brett M, Andersson J, Wager T, Poline JB. Valid conjunction inference with the minimum statistic. NeuroImage. 2005;25(3):653–660.

58. Coutanche MN, Thompson-Schil SL. Informational connectivity: Identifying synchronized discriminability of multi-voxel patterns across the brain. Frontiers in Human Neuroscience. 2013;7:36711.

